# Comparative genomics of the tardigrades *Hypsibius dujardini* and *Ramazzottius varieornatus*

**DOI:** 10.1101/112664

**Authors:** Yuki Yoshida, Georgios Koutsovoulos, Dominik R. Laetsch, Lewis Stevens, Sujai Kumar, Daiki D. Horikawa, Kyoko Ishino, Shiori Komine, Takekazu Kunieda, Masaru Tomita, Mark Blaxter, Kazuharu Arakawa

## Abstract

Tardigrada, a phylum of meiofaunal organisms, have been at the center of discussions of the evolution of Metazoa, the biology of survival in extreme environments, and the role of horizontal gene transfer in animal evolution. Tardigrada are placed as sisters to Arthropoda and Onychophora (velvet worms) in the superphylum Ecdysozoa by morphological analyses, but many molecular phylogenies fail to recover this relationship. This tension between molecular and morphological understanding may be very revealing of the mode and patterns of evolution of major groups. Similar to bdelloid rotifers, nematodes and other animals of the water film, limno-terrestrial tardigrades display extreme cryptobiotic abilities, including anhydrobiosis and cryobiosis. These extremophile behaviors challenge understanding of normal, aqueous physiology: how does a multicellular organism avoid lethal cellular collapse in the absence of liquid water? Meiofaunal species have been reported to have elevated levels of HGT events, but how important this is in evolution, and in particular in the evolution of extremophile physiology, is unclear. To address these questions, we resequenced and reassembled the genome of *Hypsibius dujardini*, a limno-terrestrial tardigrade that can undergo anhydrobiosis only after extensive pre-exposure to drying conditions, and compared it to the genome of *Ramazzottius varieornatus*, a related species with tolerance to rapid desiccation. The two species had contrasting gene expression responses to anhydrobiosis, with major transcriptional change in *H. dujardini* but limited regulation in *R. varieornatus*. We identified few horizontally transferred genes, but some of these were shown to be involved in entry into anhydrobiosis. Whole-genome molecular phylogenies supported a Tardigrada+Nematoda relationship over Tardigrada+Arthropoda, but rare genomic changes tended to support Tardigrada+Arthropoda.

## INTRODUCTION

The superphylum Ecdysozoa emerged in the Precambrian, and ecdysozoans not only dominated the early Cambrian explosion but are also dominant (in terms of species, individuals and biomass) today. The relationships of the eight phyla within Ecdysozoa remain contentious, with morphological assessments, developmental analyses and molecular phylogenetics yielding conflicting signals [1–3]. It has generally been accepted that Arthropoda, Onychophora (the velvet worms) and Tardigrada (the water bears or moss piglets) form a monophylum “Panarthropoda” [2], and that Nematoda (roundworms) are closely allied to Nematomorpha (horsehair worms). However, molecular phylogenies have frequently placed representatives of Tardigrada as sister to Nematoda [1, 3], invalidating Panarthropoda and challenging models of the evolution of complex morphological traits such as segmentation, serially repeated lateral appendages, the triradiate pharynx and a tripartite central nervous system [4, 5].

The key taxon in these disagreements is phylum Tardigrada. Nearly 1,200 species of tardigrades have been described [6]. All are members of the meiofauna-small animals that live in the water film and in interstices between sediment grains [6]. There are marine, freshwater and terrestrial species. Many species of terrestrial tardigrades are cryptobiotic: they have the ability to survive extreme environmental challenges by entering a dormant state [7]. Common to these resistances is an ability to lose or exclude the bulk of body water, and anhydrobiotic tardigrades have been shown to have tolerance to high and low temperatures (including freezing), organic solvents, X- and gamma-rays, high pressure and vacuum of space [8–15]. The physiology of anhydrobiosis in tardigrades has been explored extensively, but little is currently known about its molecular bases [16, 17]. Many other taxa have cryptobiotic abilities, including some nematodes and arthropods [18], and comparison of the mechanisms in different independent acquisitions of this trait will reveal underlying common mechanisms.

Key to developing tractable experimental models for cryptobiosis is the generation of high-quality genomic resources. Genome assemblies of two tardigrades, *Hypsibius dujardini* [19–21] and *Ramazzottius varieornatus* [22], both in the family Hypsibiidae, have been published. *H. dujardini* is a limno-terrestrial tardigrade which is easy to culture [23], while *R. varieornatus* is a terrestrial tardigrade, and highly tolerant of environmental extremes [24]. An experimental toolkit for *H. dujardini*, including RNAi and in situ hybridization is being developed [25]. *H. dujardini* is, however, poorly cryptobiotic compared to *R. varieornatus*. *H. dujardini* requires 48 hr of preconditioning at 85% relative humidity (RH) and further 24 hr in 30% RH [23] to enter cryptobiosis with high survival, while *R. varieornatus* can form a tun (the cryptobiotic form) within 30 min at 30% RH [26].

A number of anhydrobiosis-related genes have been identified in Tardigrada. Catalases, superoxide dismutases, and glutathione reductases may protect against oxidative stress [27], and chaperones, such as heat shock protein 70 (HSP70) [28–30] and others, may act to protect proteins from the denaturing effects of water loss [16, 31, 32]. Additionally, several tardigrade-specific gene families have been implicated in anhydrobiosis, based on their expression patterns. In *R. varieornatus*, cytosolic abundant heat soluble (CAHS), secretory abundant heat soluble (SAHS), late embryogenesis abundant protein mitochondrial (RvLEAM), mitochondrial abundant heat soluble protein (MAHS), and damage suppressor (Dsup) gene families have been identified [22, 33, 34]. Surprisingly, analyses of the *R. varieornatus* genome also showed extensive gene loss in the peroxisome pathway and stress signaling pathways, suggesting that this species is compromised in terms of reactive oxygen resistance and repair of cellular damage [22]. While loss of these pathways would be lethal for a normal organism, loss of these resistance pathways may be associated with anhydrobiosis.

Desiccation in some taxa induces the production of anhydroprotectants, small molecules that likely replace cellular water to stabilize cellular machinery. Trehalose, a disaccharide shown to contribute to anhydrobiosis in midges [35, 36], nematodes [37] and artemia [38], is not present in the tardigrade *Milnesium tardigradum* [31]. Coupled with the ability of *R. varieornatus* to enter anhydrobiosis rapidly (i.e. without the need for extensive preparatory biosynthesis), this suggests that tardigrade anhydrobiosis does not rely on induced synthesis of protectants. Entry into anhydrobiosis in *H. dujardini* does require active transcription during preconditioning, suggesting the activation of a genetic program to regulate physiology. Some evidence supports this inference: PPA1/2A is an inhibitor of the FOXO transcription factor, which induces anti-oxidative stress pathways, and inhibition of PPA1/2A led to high lethality in *H. dujardini* during anhydrobiosis induction [23]. As *R. varieornatus* does not require preconditioning, systems critical to anhydrobiosis in *R. varieornatus* are likely to be constitutively expressed.

*H. dujardini* and *R. varieornatus* are relatively closely related (both are members of Hypsibiidae), and both have available genome sequences. The *R. varieornatus* genome has high contiguity and scores highly in all metrics of gene completeness [22]. For *H. dujardini*, three assemblies have been published. One has low contiguity and contains a high proportion of contaminating bacterial and other sequence [19]. The other two assemblies [20, 21] eliminate most contamination, but have uncollapsed haploid segments because of unrecognized heterozygosity, estimated to be around 30~60% from k-mer distributions. The initial, low quality *H. dujardini* genome was published alongside a claim of extensive horizontal gene transfer (HGT) from bacteria and other taxa into the tardigrade genome, and a suggestion that HGT might have contributed to the evolution of cryptobiosis [19]. The “extensive” HGT claim has been robustly challenged [20, 21, 39-41], but the debate as to the contribution of HGT to cryptobiosis remains open. The genomes of these species could be exploited for understanding of rapid-desiccation versus slow-desiccation strategies in tardigrades, the importance of HGT, and the resolution of the deep structure of the Ecdysozoa. However the available genomes are not of equivalent quality.

We have generated a high quality genome assembly for *H. dujardini*, using an array of data including single-tardigrade sequencing [42] and PacBio SMRT long reads. Using a heterozygote-aware assembly method [43, 44], we minimized residual heterozygosity. Gene finding and annotation with extensive RNA-Seq data allowed us to predict a robust gene set. While most (60%) of the genes of *H. dujardini* had direct orthologues in an improved gene prediction for *R. varieornatus*, levels of synteny were very low. We identified an unremarkable proportion of potential horizontal gene transfers (maximally 1.3 – 1.8%). *H. dujardini* showed losses of peroxisome and stress signaling pathways, as described in *R. varieornatus*, as well as additional unique losses. Transcriptomic analysis of anhydrobiosis entry detected higher levels of regulation in *H. dujardini* compared to *R. varieornatus*, as predicted, including regulation of genes with anti-stress and apoptosis functions. Using single copy orthologues, we reanalyzed the position of Tardigrada within Ecdysozoa and found strong support for a Tardigrade-Nematode clade, even when data from transcriptomes of a nematomorph, onychophorans and other ecdysozoan phyla were included. However, rare genomic changes tended to support the more traditional Panarthropoda. We discuss our findings in the context of how best to improve genomics of neglected species, the biology of anhydrobiosis and the conflicting models of ecdysozoan relationships.

## RESULTS

### THE GENOME OF *H. DUJARDINI*

The genome size of *H. dujardini* has been independently estimated by densitometry to be ~ 100 Mb [20, 45], but the spans of existing assemblies exceed this, because of contamination with bacterial reads and uncollapsed heterozygosity. We generated new sequencing data (Supplementary Table 1A), including PacBio long single-molecule reads and data from single tardigrades [42], and employed an assembly strategy that eliminated evident bacterial contamination and dealt with heterozygosity. Our initial Platanus [44] genome assembly had a span of 99.3 Mb in 1,533 contigs, with an N50 length of 250 kb. Further scaffolding and gap filling [46] with PacBio reads and a Falcon [43] assembly of the PacBio reads produced a 104 Mb assembly in only 1,421 scaffolds and an N50 length of 342 kb, N90 count of 343 (Table 1). In comparison with previous assemblies, this assembly has improved contiguity and improved coverage of complete core eukaryotic genes (Complete 237, average count 1.17, Partial 240, average count 1.20) [47]. Read coverage was relatively uniform throughout the genome (Supplementary Table S2), with only a few short regions, likely repeats, with high coverage (Supplementary Figure S1).

**Table 1.** Metrics of *Hypsibius dujardini* genome assemblies.

| Data Source | This work | Edinburgh | UNC |
| --- | --- | --- | --- |
| Sequencing technologies | Illumina & PacBio | Illumina | Illumina & PacBio |
| Genome version | nHd.3.0 | nHd.2.3 | tg |
| Scaffold number | 1,421 | 13,202 | 16,175 |
| Total Scaffold Length (bp) | 104,155,103 | 134,961,902 | 212,302,995 |
| Average Scaffold Length (bp) | 73,297 | 10,222 | 13,125 |
| Longest Scaffold Length (bp) | 2,115,976 | 594,143 | 1,208,507* |
| Shortest Scaffold Length (bp) | 1,000 | 500 | 2,002 |
| N50 (bp) (no. scaffs in N50) | 342,180 (#85) | 50,531 (#701) | 17,496 (#3,422) |
| N90 (bp) (no. scaffs in N90) | 65,573 (#343) | 6,194 (#3,280) | 6,637 (#11,175) |
| CEGMA genes found (partial) | 237 (240) | 220 (241) | 221 (235) |
| CEGMA gene duplication ratio | 1.17 (1.23) | 1.35 (1.56) | 3.26 (3.53) |

\* The longest scaffolds in the tg assembly are derived from bacterial contaminants.

We identified repeats in the *H. dujardini* genome, and identified 24.2Mb (23.4%) as being repetitive elements. Simple repeats covered 5.2% of the genome, with a longest repeat unit of 8,943 bp. A more complete description of the repeat content is available in Supplementary Table S3. Seven of the eight longest repeats were of the same repeat unit (GATGGGTTTT)_n_. The long repeats of this 10 base unit were found exclusively at 9 scaffold ends and may correspond to telomeric sequence (Supplementary Table S4). The other long repeat was a simple repeat of (CAGA)_n_ and its complementary sequence (GTCT)_n_, and spanned 3.2 Mb (3% of the genome, longest repeat 5,208 bp).

We generated RNA-Seq data from active and cryptobiotic (“tun” stage) tardigrades, and developmental stages of *H. dujardini* (Supplementary Table S1B,). Gene annotation using BRAKER [48] predicted 19,901 genes, with 914 isoforms (version nHd3.0). These coding sequence predictions lacked 5’ and 3’ untranslated regions. Mapping of RNA-Seq data to the predicted coding transcriptome showed an average mapping ratio of 50%, but the mapping ratio was over 95% against the genome (Supplementary Table S5). A similar mapping pattern for RNA-Seq data to predicted transcriptome was also observed for *R. varieornatus*. Furthermore, over approximately 70% of the *H. dujardini* transcripts assembled with Trinity [49] map to the predicted transcriptome, and a larger proportion to the genome (Supplementary Table S6). RNA-seq reads that are not represented in the predicted coding transcriptome likely derived from UTR regions, unspliced introns or from promiscuous transcription. We were able to infer functional and similarity annotations for ~50% of the predicted proteome (Table 2).

**Table 2.**
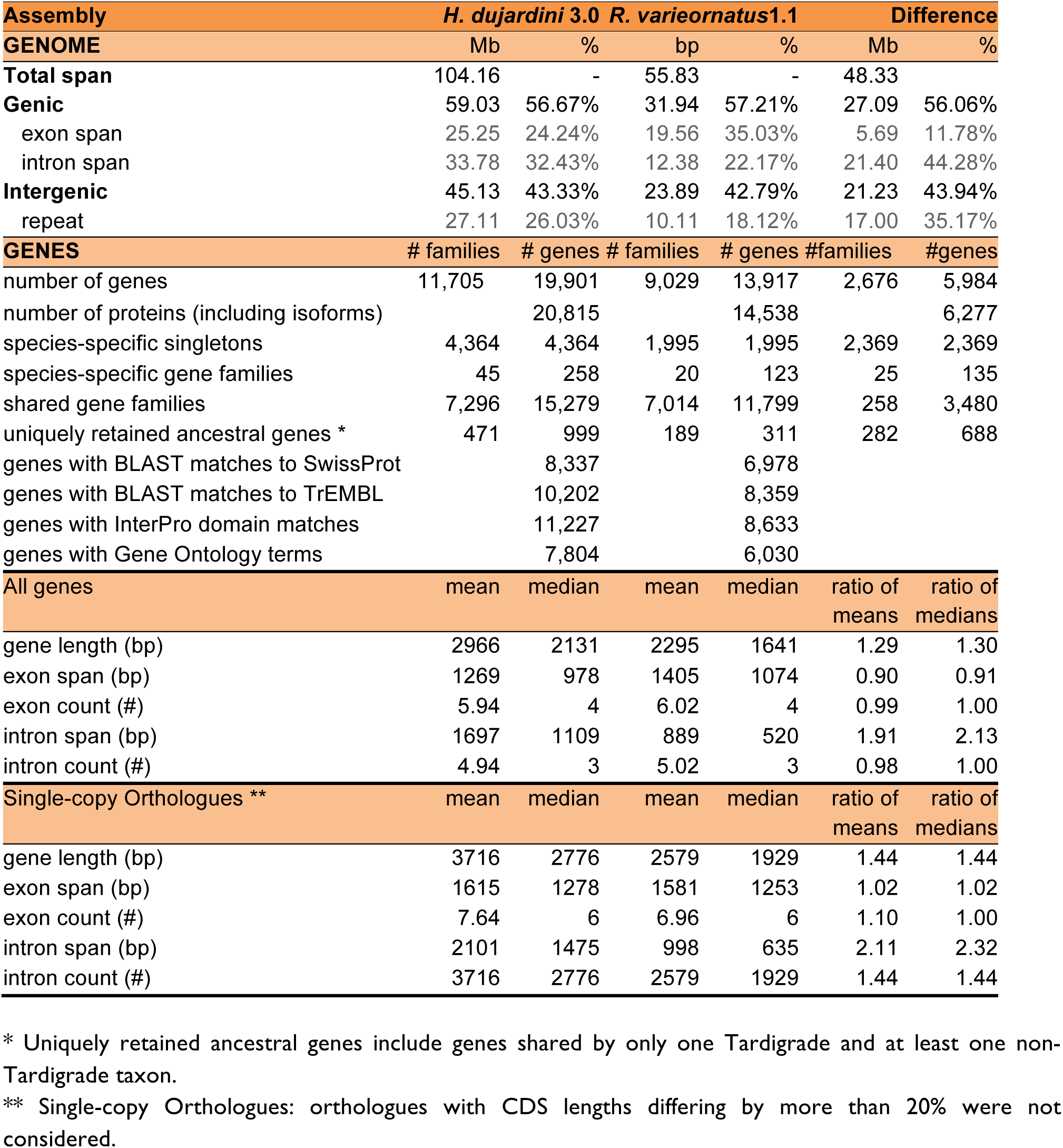
Comparison of the genomes of *H. dujardini* and *R. varieornatus*.

| Assembly | *H. dujardini* 3.0 |  | *R. varieornatus*1.1 |  | Difference |  |
| --- | --- | --- | --- | --- | --- | --- |
| GENOME | Mb | % | bp | % | Mb | % |
| **Total span** | 104.16 | - | 55.83 | - | 48.33 |  |
| **Genic** | 59.03 | 56.67% | 31.94 | 57.21% | 27.09 | 56.06% |
| exon span | 25.25 | 24.24% | 19.56 | 35.03% | 5.69 | 11.78% |
| intron span | 33.78 | 32.43% | 12.38 | 22.17% | 21.40 | 44.28% |
| **Intergenic** | 45.13 | 43.33% | 23.89 | 42.79% | 21.23 | 43.94% |
| repeat | 27.11 | 26.03% | 10.11 | 18.12% | 17.00 | 35.17% |
| GENES | # families | # genes | # families | # genes | #families | #genes |
| number of genes | 11,705 | 19,901 | 9,029 | 13,917 | 2,676 | 5,984 |
| number of proteins (including isoforms) |  | 20,815 |  | 14,538 |  | 6,277 |
| species-specific singletons | 4,364 | 4,364 | 1,995 | 1,995 | 2,369 | 2,369 |
| species-specific gene families | 45 | 258 | 20 | 123 | 25 | 135 |
| shared gene families | 7,296 | 15,279 | 7,014 | 11,799 | 258 | 3,480 |
| uniquely retained ancestral genes * | 471 | 999 | 189 | 311 | 282 | 688 |
| genes with BLAST matches to SwissProt |  | 8,337 |  | 6,978 |  |  |
| genes with BLAST matches to TrEMBL |  | 10,202 |  | 8,359 |  |  |
| genes with InterPro domain matches |  | 11,227 |  | 8,633 |  |  |
| genes with Gene Ontology terms |  | 7,804 |  | 6,030 |  |  |
| All genes | mean | median | mean | median | ratio of means | ratio of medians |
| gene length (bp) | 2966 | 2131 | 2295 | 1641 | 1.29 | 1.30 |
| exon span (bp) | 1269 | 978 | 1405 | 1074 | 0.90 | 0.91 |
| exon count (#) | 5.94 | 4 | 6.02 | 4 | 0.99 | 1.00 |
| intron span (bp) | 1697 | 1109 | 889 | 520 | 1.91 | 2.13 |
| intron count (#) | 4.94 | 3 | 5.02 | 3 | 0.98 | 1.00 |
| Single-copy Orthologues ** | mean | median | mean | median | ratio of means | ratio of medians |
| gene length (bp) | 3716 | 2776 | 2579 | 1929 | 1.44 | 1.44 |
| exon span (bp) | 1615 | 1278 | 1581 | 1253 | 1.02 | 1.02 |
| exon count (#) | 7.64 | 6 | 6.96 | 6 | 1.10 | 1.00 |
| intron span (bp) | 2101 | 1475 | 998 | 635 | 2.11 | 2.32 |
| intron count (#) | 3716 | 2776 | 2579 | 1929 | 1.44 | 1.44 |

\* Uniquely retained ancestral genes include genes shared by only one Tardigrade and at least one non- Tardigrade taxon. \*\* Single-copy Orthologues: orthologues with CDS lengths differing by more than 20% were not considered.

We identified eighty-one 5.8S rRNA, two 18S rRNA, and three 28S rRNA loci with RNAmmer [50]. Scaffold0021 contains both 18S and 28S loci, and it is likely that multiple copies of the ribosomal RNA repeat locus have been collapsed in this scaffold, as it has very high read coverage (~5,400 fold, compared to ~113x fold overall, suggesting ~48 copies). tRNAs for each amino acid were found (Supplementary Figure S2) [51]. Analysis of miRNA-Seq data with miRDeep [52] predicted 507 mature miRNA loci (Supplementary Data S1), of which 185 showed similarity with sequences in miRbase [53].

The *H. dujardini* nHd.3.0 genome assembly is available on a dedicated ENSEMBL [54] server, http://ensembl.tardigrades.org, where it can be compared with previous assemblies of *H. dujardini* and with the *R. varieornatus* assembly. The ENSEMBL database interface includes an application-programming interface (API) for scripted querying [55]. All data files (including supplementary data files and other analyses) are available from http://download.tardigrades.org, and a dedicated BLAST server is available at http://blast.tardigrades.org. All raw data files have been deposited in INSDC databases (NCBI and SRA, Supplementary Table SIA-C) and the assembly with a slightly curated annotation (nHd3.l) has been submitted to NCBI under the accession ID MTYJ00000000.

### COMPARISONS WITH *RAMAZZOTTIUS VARIEORNATUS*

We compared this high-quality assembly of *H. dujardini* to that of *R. varieornatus* [22]. In initial comparisons we noted that *R. varieornatus* had a high proportion of single-exon loci that had no *H. dujardini* (or other) homologues. Reasoning that this might be a technical artifact due to the different gene finding strategies used, we updated gene models for *R. varieornatus* using the BRAKER pipeline [48] with additional comprehensive RNA-Seq of developmental stages (Supplementary Table 1C). The new prediction includes 13,917 protein-coding genes (612 isoforms). This lower gene count compared to the original (19,521 genes) is largely driven by a reduction in single-exon genes that had no transcript support: from 5,626 in version 1 to 1,777 in the current annotation. Comparing the two gene sets, 12,752 of the BRAKER-predicted genes were also found in the original gene set. In both species, some of the predicted genes may derive from transposons, as 2,474 *H. dujardini* and 626 *R. varieornatus* proteins match Dfam domains [56]. While several of these putatively transposon-derived predictions have a Swiss-Prot [57] homologue *(H. dujardini* 915, 36.98%, *R. varieornatus*:274, 43.76%), a large proportion had very low expression levels. In *H. dujardini*, 71.6% (1551 loci) had less than 10 transcripts per million mapped fragments (TPM) and 46.2% (1144) less than 1 TPM. In *R. varieornatus* 67.57% (423) had less than 10 and 37.9% (237) less than 1 TPM. Among the gene models of *H. dujardini*, 39 were predicted to contain 4 “ochre” (TAA), 12 “amber” (TAG), and 23 “opal” (TGA) stop codons.

One striking difference between the two species was in their genome size, as represented by assembly span: the *R. varieornatus* assembly had a span of 55 Mb, half that of *H. dujardini* (Table 2). This difference could have arisen through whole genome duplication, segmental duplication, or more piecemeal processes of genome expansion or contraction between the two species. *H. dujardini* had 5,984 more predicted genes than *R. varieornatus*. These spanned ~23 Mb, and thus accounted for about half of the additional span. We observed no major difference in number of exons between orthologues or in the predicted gene set as a whole. However, comparing orthologues, the intron span per gene in *H. dujardini* was on average twice that in *R. varieornatus* (Figure 1B), and the gene length (measured as start codon to stop codon in the coding exons) was ~1.3 fold longer in *H. dujardini* (Supplementary Figure S3). The non-genic component of noncoding DNA was greater in *H. dujardini*, and this increase was largely explained by an increase in the repeat content (27 Mb in *H. dujardini, versus* 10 Mb in *R. varieornatus)*.

**Figure 1.**
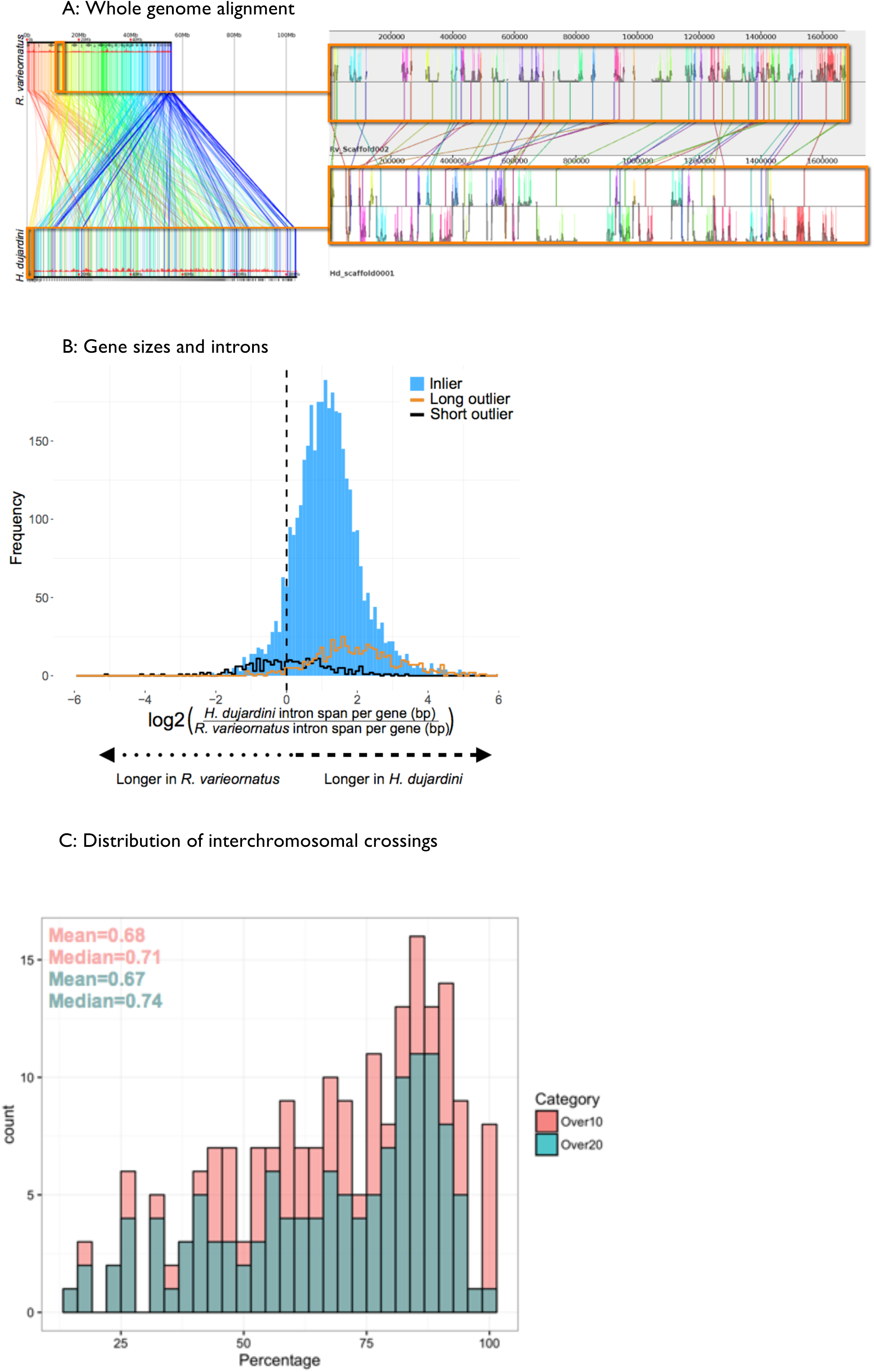
The genomes of *Hypsibius dujardini* and *Ramazzottius varieornatus*. **A Linkage conservation but limited synteny between *H. dujardini* and *R. varieornatus***. Whole genome alignment was performed with Murasaki (reference) The left panel shows the whole genome alignment. Similar regions are linked by a line colored following a spectrum based on the start position in *R. varieornatus*. To the right is a re-alignment of the initial segment of *H. dujardini* scaffold0001 (lower), showing matches to several portions of *R. varieornatus* scaffold0002 (above), illustrating the several inversions that must have taken place. The histograms show pairwise nucleotide sequence identity between these two segments. **B Gene sizes and introns**. *H. dujardini* genes are longer because of expanded introns. Frequency histogram of log2 ratio of intron span per gene in *H. dujardini* compared to *R. varieornatus*. **C Distribution of interchromosomal crossings**. To test conservation of gene neighbourhood conservation, we asked whether genes found together in *H. dujardini* were also found close together in *R. varieornatus*. Taking the set of genes on each long *H. dujardini* scaffold, we identified the locations of the reciprocal best BLAST hit orthologues in *R. varieornatus*, and counted the maximal proportion mapping to one *R. varieornatus* scaffold. *H. dujardini* scaffolds were were binned and counted by this proportion. As short scaffolds, with fewer genes, might bias this analysis, we performed analyses independently on scaffolds with > 10 genes and scaffolds with >20 genes.

Whole genome alignments of *R. varieornatus* and *H. dujardini* using Murasaki [58] revealed a low level of synteny but evidence for conserved linkage at the genome scale, with little conservation of gene order beyond a few loci. For example, comparison of Scaffold002 of *R. varieornatus* with scaffold0001 of *H. dujardini* showed long-range linkage, with many shared (genome-wide bidirectional best BLAST hit) loci spread across ~1.7 Mb of the *H. dujardini* genome (Figure 1A). Furthermore, we found that a high proportion of genes located on the same scaffold in *H. dujardini* were located in the same scaffold in *R. varieornatus* as well, implying that interchromosomal rearrangement may be the reason for the low level of synteny (Figure 1C).

We clustered the *H. dujardini* and new *R. varieornatus* proteomes with a selection of other ecdysozoan and outgroup lophotrochozoan proteomes (Supplementary Table S7) using OrthoFinder [59], including proteomes from fully-sequenced genomes and proteomes derived from (likely partial) transcriptomes in independent analyses (Supplementary table S11). These protein clusters were used for subsequent identification of orthologues for phylogenetic analysis, and for patterns of gene family expansion and contraction, using kinfin [60].

Orthologue clustering for the analysis of gene families (see Supplementary table S7 for datasets used) generated 144,610 clusters composed of 537,608 proteins (spanning 210,412,426 aminoacids) from the 29 selected species. Of these clusters, 87.9% are species specific (with singletons accounting for 11.6% of amino acid span, and multi-protein clusters accounting for 1.2% of span). While only 12.1% of clusters contain members from two or more proteomes, these account for the majority of amino acid span (87.2%). *H. dujardini* had more species-specific genes than did *R. varieornatus*, and had more duplicate genes in gene families shared with *R. varieornatus* (Table 2). *H. dujardini* also had more genes shared with non-tardigrade outgroups, suggesting loss in *R. varieornatus*.

One hundred and fifteen clusters had more members in tardigrades compared to the other taxa, and three had fewer members, based on uncorrected Mann-Whitney U-test probabilities <0.01, but no clusters had differential presence when the analyses were Bonferroni corrected (see Supplementary Data S8: Tardigrade_counts_representation_tests). In nine of the clusters with tardigrade overrepresentation, the tardigrades had more than four times as many members as the average of the other species. Cluster OG0000104 had 284 members, 276 of which derive from the tardigrades. This cluster was annotated as having a receptor ligand binding domain, but was not otherwise distinguished. OG004022 had 5 members in *H. dujardini* and 9 members in *R. variornatus*, but a mode of 1 (and maximum of 4 in *Octopus bimaculatus*) in other species. It is a member of a deeply conserved, otherwise uncharacterized, transmembrane protein family of unknown function. OG0001636 gathers a deeply conserved ATP-binding cassette family, and *while the 27 other species had a mode of 1 (and a maximum of 2), R. varieornatus had 4 and H. dujardini* had 9 copies. OG0002927 encodes protein kinases, present in 23 of the 29 species, with 6 in *H. dujardini*, 5 in *R. varieornatus* and a mode of 1 elsewhere. OG0004228 is annotated as a relish-like transcription factor, and has 1 copy in the non-tardigrade species (except for two insects with 2) and 5 copies in each tardigrade. OG0001359, with 1 copy in most species, 8 in *H. dujardini*, 8 in *R. varieornatus*, and 4 in *Solenopsis invictus*, is likely to be a SAM-dependent methyltransferase (type 11), possibly involved in coenzyme biosynthesis. OG001949 had 1 copy in most species but 6 in *H. dujardini* and 4 in *R. varieornatus*, and is annotated as a RAB GTP hydrolase. OG0003870 was unannotated (containing only matches to domain of unknown function DUF1151), and elevated in *R. varieornatus* (9 copies) compared to other species (mode of 1; *H. dujardini* had 2). The three clusters with depletion in the tardigrades were OG0000604, encoding an exoribonuclease (1 copy in each tardigrade, but an average of three copies in the other 27 species), OG0000950, a 3'5'-cyclic nucleotide phosphodiesterase (1 in tardigrades versus 2.3 elsewhere) and OG00001138, an EF-hand protein (1 in tardigrades versus 2 elsewhere).

### HORIZONTAL GENE TRANSFER IN TARDIGRADE GENOMES

HGT is an interesting but contested phenomenon in animals. Many newly sequenced genomes have been reported to have relatively high levels of HGT, and genomes subject to intense curation efforts tend to have lower HGT estimates. We performed *ab initio* gene finding on the genomes of the model species *Caenorhabditis elegans* and *Drosophila melanogaster* with Augustus [61] and used the HGT index approach [62], which simply classifies loci based on the ratio of their best BLAST scores to ingroup and potential donor taxon databases, to identify candidates. Compared with their mature annotations, we found elevated proportions of putative HGTs in both species (*C*. *elegans*: 2.09% of all genes, *D. melanogaster:* 4.67%). We observed similarly elevated rates of putative HGT loci, as assessed by the HGT index, in gene sets generated by *ab inito* gene finding in additional arthropod and nematode genomes compared to their mature annotations (Figure 2A, Supplementary Table S8). Thus the numbers of HGT events found in the genomes of *H. dujardini* and *R. varieornatus* are likely to be overestimated, even after sequence contamination has been removed.

**Figure 2.**
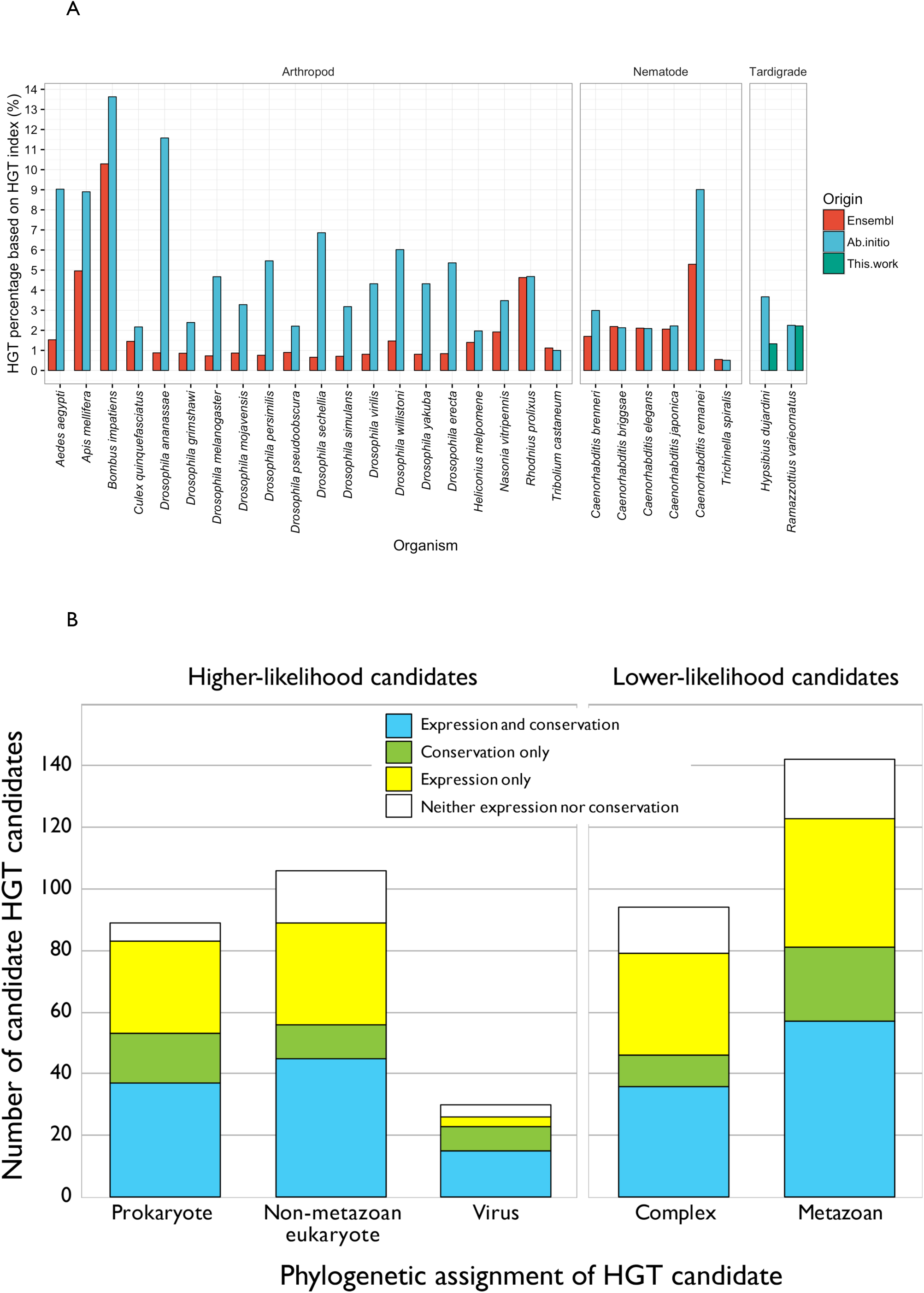
Horizontal gene transfer in *Hypsibius dujardin*. **A Horizontal gene transfer in *Hypsibius dujardini***. For a set of assembled arthropod and nematode genomes, genes were re-predicted *ab initio* with Augustus. Putative HGT loci were identified using the HGT index for the longest transcript for each gene from the new and the ENSEMBL reference gene sets. In most species the *ab initio* gene sets had elevated numbers of potential HGT loci compared to their ENSEMBL representations. **B Classification of HGT candidates in *H. dujardini***. Classification of the initial HGT candidates identified in *H. dujardini* by their phylogenetic annotation (Prokaryote, non-metazoan Eukaryote, virus, metazoan and complex), their support in RNA-Seq expression data and the presence of a homologue in *R. varieornatus*.

Using the HGT index approach we identified 463 genes (2.32% of all genes) as potential HGT candidates in *H. dujardini*. Using Diamond BLASTX [63] instead of standard BLASTX [64], made only a minor difference in the number of potential HGT events predicted (446 genes, 2.24%). We sifted the initial 463 *H. dujardini* candidates through a series of biological filters. A true HGT locus will show affinity with its source taxon when analyzed phylogenetically (a more sensitive test than simple BLAST score ratio). Just under half of the loci (225) were confirmed as HGT events by RAxML [65] analysis of aligned sequences (Figure 2B). HGT genes are expected to be incorporated into the host genome and to persist through evolutionary time. Only 214 of the *H. dujardini* candidates had homologues in *R. varieornatus*, indicating phylogenetic perdurance (Supplementary Data S2). Of these shared candidates, 113 were affirmed by phylogeny. HGT loci will acquire gene structure and expression characteristics of their host, metazoan genome. One third (168) of the HGT candidates had RNA-Seq expression values at or above the average for all genes. While metazoan genes usually contain spliceosomal introns, and 367 of the candidate HGT gene models included introns, we regard this a lower-quality validation criterion as gene finding algorithms are programmed to identify introns. Therefore our minimal current estimate for HGT into the genome of *H. dujardini* is 113 genes (0.57%, out of 19,901 loci) and the upper bound is 463 (2.33%). This is congruent with estimates of 1.58% HGT candidates (out of 13,917 genes) for *R. varieornatus* [22].

The putative HGT loci tended to be clustered in the tardigrade genomes, with gene neighbors of HGT loci also predicted to be HGT. We found 58 clusters of HGT loci in *H. dujardini*, and 14 in *R. varieornatus* (Supplementary Figure S4). Several of these gene clusters were comprised of orthologous genes, and may have arisen through tandem duplication. The largest clusters included over 5 genes from the same gene family (Supplementary Data S3).

### THE GENOMICS OF ANHYDROBIOSIS IN TARDIGRADES

We explored the *H. dujardini* proteome and the reannotated *R. varieornatus* proteome for loci implicated in anhydrobiosis. The anhydrobiosis machinery in *R. varieornatus* includes members of the cytosolic abundant heat soluble (CAHS), secretory abundant heat soluble (SAHS), late embryogenesis abundant protein mitochondrial (RvLEAM), mitochondrial abundant heat soluble protein (MAHS), and damage suppressor (Dsup) families [22, 33, 34]. In the new *R. varieornatus* proteome, we found 16 CAHS loci and 13 SAHS loci and one copy each of MAHS, RvLEAM and Dsup. In *H. dujardini* we identified 12 CAHS loci, 10 SAHS loci and single members of the RvLEAM and MAHS families (Supplementary Table S9). Direct interrogation of the *H. dujardini* genome with *R. varieornatus* loci identified an additional possible CAHS-like locus and an additional SAHS-like locus. We found no evidence for a *H. dujardini* homologue of Dsup. Phylogenetic analyses revealed a unique duplication of CAHS3 in *R. varieornatus*. No SAHS2 ortholog was found in *H. dujardini* (Supplementary Figure S5), and most of the *H. dujardini* SAHS loci belonged to a species-specific expansion that was orthologous to a single *R. varieornatus* SAHS locus, RvSAHS13. SAHS1-like genes in *H. dujardini* and SAHS1- and SAHS2-like genes in *R. varieornatus* were locally duplicated, forming SAHS clusters on single scaffolds.

*R. varieornatus* was reported to have undergone extensive gene loss in the stress responsive transducer of CREB protein mTORC and in the peroxisome pathway, which generates H_2_O_2_ during the beta-oxidation of fatty lipids. *H. dujardini* was similarly compromised (Figure 3A). We identified additional gene losses in the peroxisome pathway in *H. dujardini:* peroxisome proteins PEK5, PEK10, and PEK12, while present in *R. varieornatus*, were not found in *H. dujardini*.

**Figure 3.**
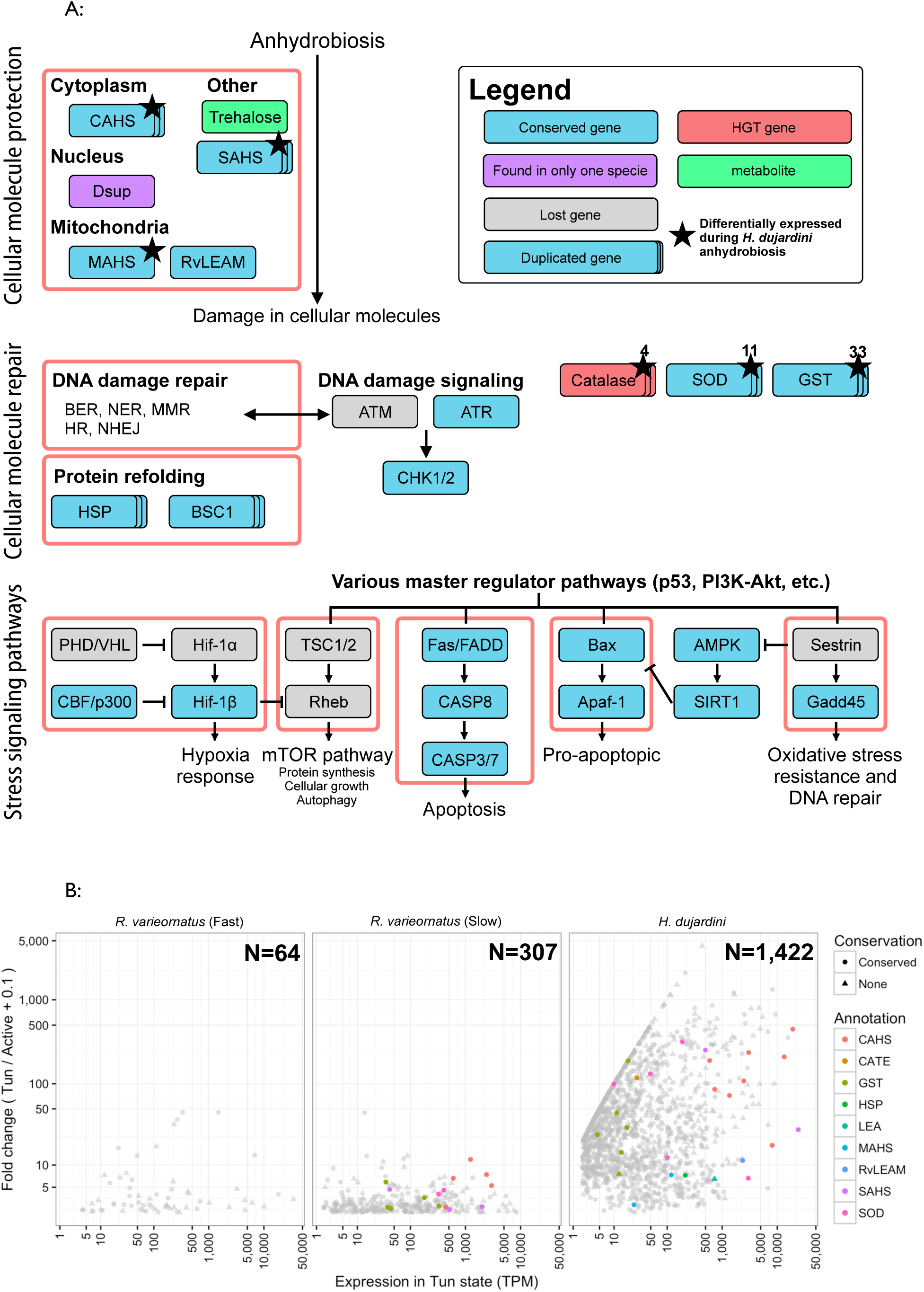
The genomics of anhydrobiosis in tardigrades. **A Gene losses in Hypsibiidae**. Gene losses were detected by mapping to KEGG pathways using KAAS, and validated by BLAST v2.2.22 TBLASTN search of KEGG ortholog gene amino acid sequences. Light blue and gray boxes indicate genes conserved and lost in both tardigrades, respectively. Furthermore, purple boxes represent genes retained in only one species, and red as genes that have been detected as HGT. Numbers on the top right of boxes indicates copy numbers of multiple copy genes in *H. dujardini*. Genes annotated as CASP3 and CDC25A have contradicting annotation with KAAS and Swiss-Prot, however the KAAS annotation was used. **B Differential gene expression in tardigrades on entry to the anhydrobiotic state.** The TPM expression for each samples were calculated with Kallisto, and the fold change between active and tun, and the TPM expression in the Tun state were plotted. Genes that likely contribute to anhydrobiosis were colored, and genes that had an ortholog in the other species were shaped as a circle.

Three of the orthology clustering families had uncorrected (Mann-Whitney U-test, p<0.01) overrepresentation in the tardigrades and contained members that were differentially expressed during anhydrobiosis. Cluster OG000684 has members from 28 of the 29 species, but *H. dujardini* and *R. varieornatus* had more copies (33 and 8, respectively) than any other (mode of 1 and mean of 1.46 copies per species, with a maximum of 4 in the moth *Plutella xylostella*). Proteins in the cluster were annotated with domains associated with ciliar function. OG0002660 contained three proteins from each of *H. dujardini* and *R. varieornatus*, but a mean of 1.2 from the other species. OG0002660 was annotated as fumarylacetoacetase, which acts in phenyalanine metabolism. Fumarylacetoacetase has been identified as a target of SKN-1 induced stress responses in *C. elegans* [66]. OG0002103 was also overrepresented in the tardigrades (3 in each species), while 23 of the other species had 1 copy. Interestingly the extremophile nematode *Plectus murrayi* had 4 copies. OG0002103 was annotated as GTP cyclohydrolase, involved in formic acid metabolism, including tetrahydrobioterin synthesis. Tetrahydrobioterin is a cofactor of aromatic amino acid hydroxylases, which metabolise phenylalanine.

We used orthology clustering to explore family sizes of genes with functions associated with anhydrobiosisz (Supplementary Data S5: Tardigrade_DEGs.functional_annotation). Proteins with functions related to protection from oxidants, such as superoxide dismutase (SOD) and peroxiredoxin, were found to have been extensively duplicated in tardigrades. In addition, the mitochondrial chaperone (BSC1), osmotic stress related transcription factor NFAT5, and apoptosis related gene PARP families were expanded in tardigrades. Chaperones were extensively expanded in *H. dujardini* (HSP70, DnaK, and DnaJ subfamily C-5, C-13, B-12), and the DnaJ subfamily B3, B-8 was expanded in *R. varieornatus*. In *H. dujardini*, we found five copies of DNA repair endonuclease XPF, which functions in the nucleotide-excision repair pathway, and in *R. varieornatus*, four copies of the double-stranded break repair protein MRE11 (as reported previously [22]) and additional copies of DNA ligase 4, from the non-homologous end joining pathway.

In both *R. varieornatus* [22] and *H. dujardini* some of the genes with anhydrobiosis related function appear to have been acquired through HGT. All copies of catalase were high-confidence HGTs. *R. varieornatus* had eleven copies of trehalase (nine trehalases and two acid trehalase-like proteins). Furthermore, we found that *H. dujardini* does not have an ortholog of trehalose phosphatase synthase, a gene required for trehalose synthesis, however *R. varieornatus* has a HGT derived homolog (Supplementary Figure S6A). The ascorbate synthesis pathway appears to have been acquired through HGT in *H. dujardini*, and a horizontally acquired L-gulonolactone oxidase was identified in *R. varieornatus* (Supplementary figure S6B).

To identify gene functions associated with anhydrobiosis, we explored differential gene expression in both species in fully hydrated and post-desiccation samples from both species. We compared the single individual RNA-Seq of *H. dujardini* undergoing anhydrobiosis [42] with new data for *R. varieornatus* induced to enter anhydrobiosis in two ways: slow desiccation ~24 hr) and fast desiccation (~30 min). Successful anhydrobiosis was assumed when >90% of the samples prepared in the same chamber recovered after rehydration. Many more genes (1,422 genes, 7.1%) were differentially upregulated by entry into anhydrobiosis in *H. dujardini* than in *R. varieornatus* (64 genes, 0.5%, in fast desiccation and 307 genes, 2.2%, in slow desiccation, Supplementary Data File S5). The fold change distribution of the whole transcriptome of *H. dujardini* (8.33/0.910±69.90) was significantly broader than those of both fast (0.67/0.4758±2.25) and slow (0.77/0.6547±0.79) desiccation *R. varieornatus* (U-test, p-value <0.001, mean/median±S.D., Figure 3B).

Several protection-related genes were differentially expressed in anhydrobiotic *H. dujardini*, including CAHS (8 loci of 15), SAHS (2 of 10), RvLEAM (1 of 1), and MAHS (1 of 1). Loci involved in reactive oxygen protection (5 superoxide dismutase genes, 6 glutathione-S transferase genes, and a catalase gene, 1 LEA gene) were upregulated under desiccation. Interestingly, two trehalase loci were upregulated, even though we were unable to identify a trehalose-6-phosphate synthase (TPS) locus in *H. dujardini*. In addition to these effector loci, we identified differentially expressed transcription factors that may regulate anhydrobiotic responses. Similarly, two calcium-signaling inhibitors, calmodulin (CaM) and a cyclic nucleotide gated channel (CNG-3), were both upregulated, which may drive cAMP synthesis through adenylate cyclase.

Although *R. varieornatus* is capable of rapid anhydrobiosis induction, complete desiccation is unlikely to be as rapid in natural environments, and regulation of gene expression under slow desiccation might reflect a more likely scenario. Fitting this expectation, five CAHS loci and a single SAHS locus were upregulated after slow desiccation, but none were differentially expressed following rapid desiccation. Most *R. varieornatus* CAHS and SAHS orthologs had high expression in the active state, several over 1,000 TPM. In contrast, *H. dujardini* CAHS and SAHS orthologs had low resting expression (median 0.7 TPM), and were upregulated (median 1916.8 TPM) on anhydrobiosis induction. The contributions to anhydrobiosis of additional genes identified as upregulated (including cytochrome P450, several solute carrier families, and apolipoproteins) remains unknown.

Some genes differentially expressed in both *H. dujardini* and *R. varieornatus* slow-desiccation anhydrobiosis were homologous. Of the 1,422 DEGs from *H. dujardini*, 121 genes were members of 70 protein families that also contained 115 *R. varieornatus* DEGs. These included CAHS, SAHS, GST, and SOD gene families, but in each case *H. dujardini* had more differentially expressed copies compared to *R. varieornatus*. Annotations of other gene families included metalloproteinase, calcium binding receptor and G-protein coupled receptor, suggesting that these functions may participate in cellular signaling during induction of anhydrobiosis. Many more (887) gene families included members that were upregulated by anhydrobiosis in *H. dujardini*, but unaffected by desiccation in *R. varieornatus*. These gene families included 1,879 *R. varieornatus* genes, some (154) were highly expressed in the active state (TPM >100).

### PHYLOGENETIC RELATIONSHIPS OF TARDIGRADA

We inferred orthologous gene clusters between *H. dujardini, R. varieornatus*, a set of taxa from other ecdysozoan phyla, and two lophotrochozoan outgroup taxa. We analyzed one dataset that contained only taxa with whole genome data, and a second that also included taxa with transcriptome data. The second dataset had wider representation of phyla, including Nematomorpha, Onychophora and Kinorhyncha. After careful scrutiny of putative orthologous gene clusters to eliminate evident paralogous sequences, alignments from selected loci were concatenated into a supermatrix. The genome supermatrix was trimmed to remove low-quality alignment regions. It included 322 loci from 28 taxa spanning 67,256 aligned residues, and had 12.5% missing data. The genome+transcriptome supermatrix was not trimmed; included 71 loci from 37 taxa spanning 68,211 aligned residues, and had 27% missing data. Phylogenomic analyses were carried out in RAxML (using the General Time Reversible model with Gamma distribution of rates model, GTR+G) and PhyloBayes (using a GTR plus rate categories model, GTR-CAT+G).

The genome phylogeny (Figure 4A) strongly supported Tardigrada as a sister to monophyletic Nematoda. Within Nematoda and Arthropoda, the relationships of species were as found in previous analyses, and the earliest branching taxon in Ecdysozoa was Priapulida. RAxML bootstrap and PhyloBayes bayes proportion support was high across the tree, with only two internal nodes in Nematoda and Arthropoda receiving less-than maximal support. Analysis of RAxML phylogenies derived from the 322 individual loci revealed a degradation of support deeper in the tree, with 53% of trees supporting a monophyletic Arthropoda, 56% supporting Tardigrada plus Nematoda, and 54% supporting the monophyly of Arthropoda plus Tardigrada plus Nematoda (Figure 4A). The phylogeny derived from the genome+transcriptome dataset (Figure 4B) also recovered credibly resolved Nematoda and Arthropoda, and, as expected, placed Nematomorpha as sister to Nematoda. Tardigrada was again recovered as sister to Nematoda plus Nematomorpha, with maximal support. Priapulida plus Kinorhyncha was found to arise basally in Ecdysozoa. Unexpectedly, Onychophora, represented by three transcriptome datasets, was sister to an Arthropoda plus (Tardigrada, Nematomorpha, Nematoda) clade, again with high support.

**Figure 4.**
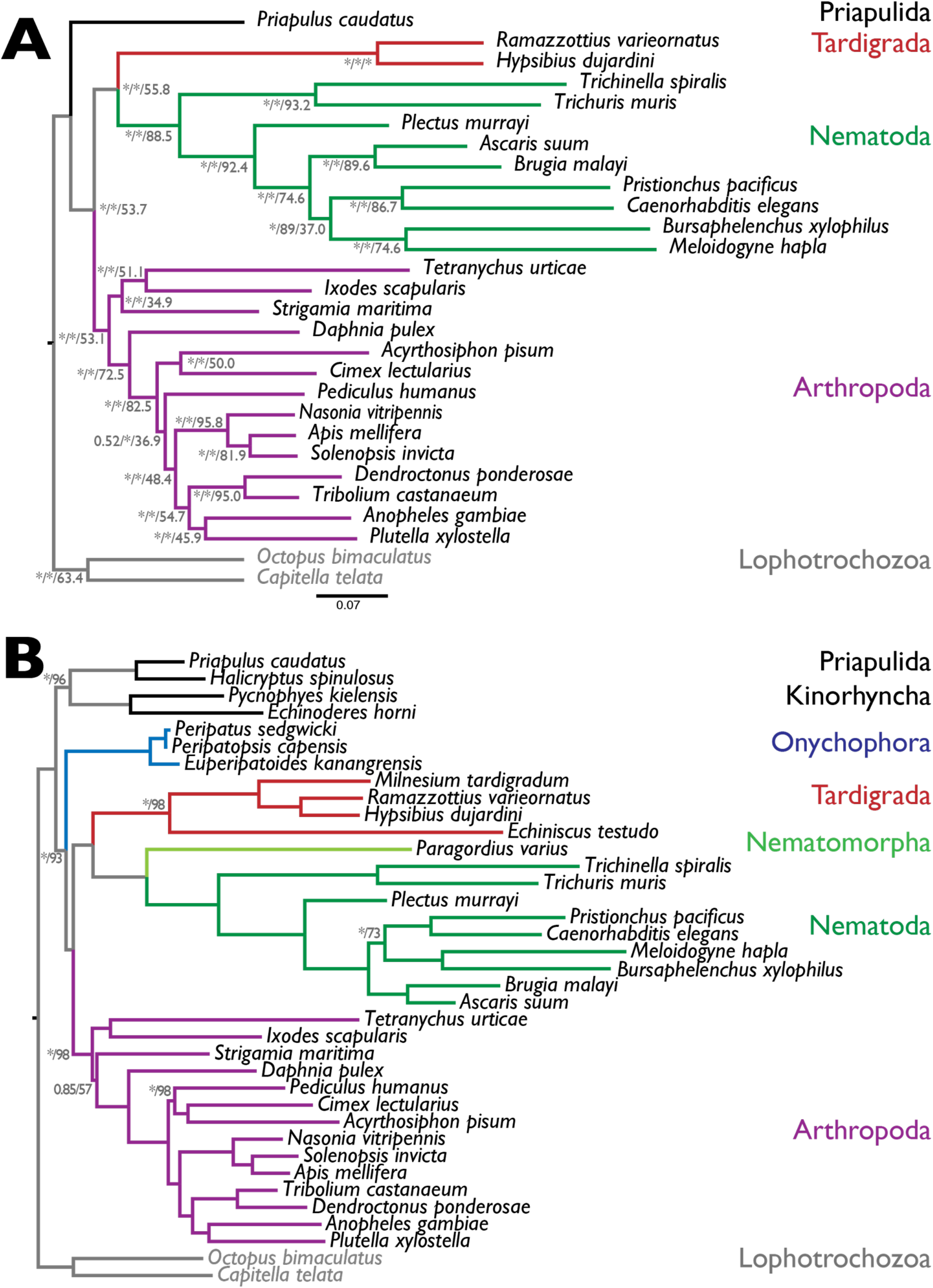
Phylogeny of Ecdysozoa. **A** Phylogeny of 28 species from 5 phyla, based on 322 loci derived from whole genome sequences, and rooted with the lophotrochozoan outgroup. Labels on nodes are Bayes proportions from PhyloBayes analysis / bootstrap proportions from RAxML maximum likelihood bootstraps / proportion of trees of individual loci supporting each bipartition. Note that different numbers of trees were assessed at each node, depending on representation of the taxa at each locus. * indicates maximal support (Bayes proportion of 1.0 or RAxML bootstrap of 1.0). **B** Phylogeny of 36 species from 8 phyla, based on 71 loci derived using PhyloBayes from whole genome and transcriptome sequences, and rooted with the lophotrochozoan outgroup. All nodes had maximal support in Bayes proportions and RAxML bootstrap, except those labeled (Bayes proportion, ^*^= 1.0 / RAxML bootstrap).

### RARE GENOMIC CHANGES AND TARDIGRADE RELATIONSHIPS

Rare genomic changes can be used as strong parsimony markers of phylogenetic relationships that are hard to resolve using model-based sequence analyses. An event shared by two taxa can be considered to support their relationship where the likelihood of the event is *a priori* expected to be vanishingly small. We tested support for a Nematoda-Tardigrada clade in rare changes in core developmental gene sets and protein family evolution.

HOX genes are involved in anterior-posterior patterning across the Metazoa, with a characteristic set of paralogous genes present in most animal genomes, organized as a tightly regulated cluster. We surveyed HOX loci in tardigrades and relatives (Figure 5A). The ancestral cluster is hypothesized to have included HOX1, HOX2, HOX3, HOX4, HOX5, and a HOX6-8 like locus and HOX9. The HOX6-8 and HOX9 types have undergone frequent, independent expansion and contraction during evolution, and HOX clustering has broken down in some species. HOX complements are generally conserved between related taxa, and gain and loss of HOX loci can be considered a rare genomic change. In the priapulid *Priapulus caudatus* nine HOX loci have been described [67], but no HOX6-8/AbdA-like gene was identified. All arthropods surveyed (including representatives of the four classes) had a complement of HOX loci very similar to that of *D. melanogaster*, with at least ten loci including HOX6-8 and HOX9. Some HOX loci in some species have undergone duplication, particularly HOX3/zen. In the mite *Tetranychus urticae* and the salmon louse *Lepeoptheirius salmonis* we identified “missing” HOX genes in the genome. For Onychophora, the sister group to Arthropoda, HOX loci have only been identified through PCR screens [68, 69], but they appear to have the same complement as Arthropoda.

**Figure 5.**
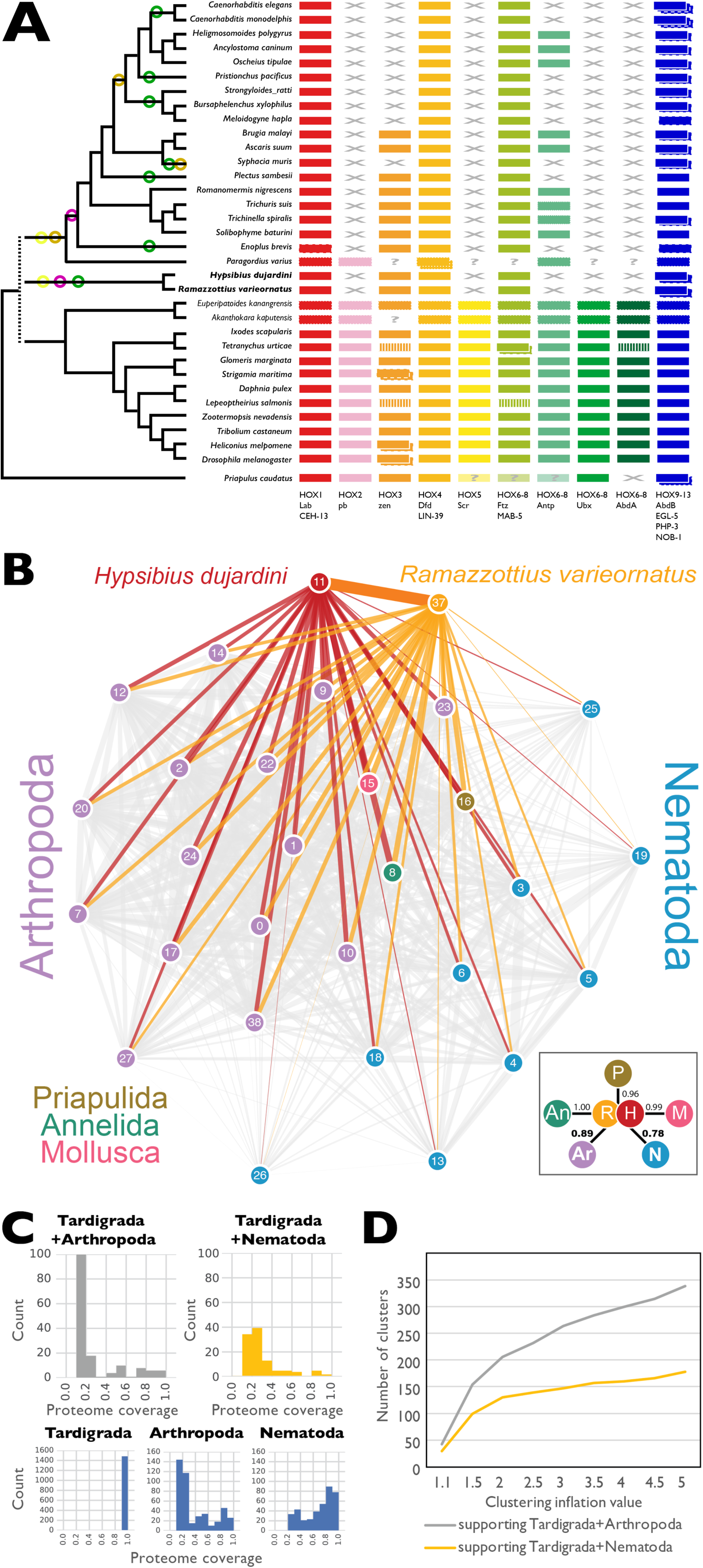
The position of tardigrada in ecdysozoa. **A HOX genes in tardigrades and other Ecdysozoa.** HOX gene losses in Tardigrada and Nematoda. HOX gene catalogues of tardigrades and other Ecdysozoa were collated by screening ENSEMBL Genomes and WormBase Parasite. HOX orthology groups are indicated by different colours. Some “missing” HOX loci were identified by BLAST search of target genomes (indicated by vertical striping of the affected HOX). “?” indicates that presence/absence could not be confirmed because the species was surveyed by PCR or transcriptomics; loci identified by PCR or transcriptomics are indicated by a dotted outline. “X” indicates that orthologous HOX loci were not present in the genome of that species. Some species have duplications of loci mapping to one HOX group, and these are indicated by boxes with dashed outlines. The relationships of the species are indicated by the cladogram to the left, and circles on this cladogram indicate Dollo parsimony mapping of events of HOX group loss on this cladogram. Circles are coloured congruently with the HOX loci. **B,C,D Evolution of gene families under different hypotheses of tardigrade relationships.** **B** Tardigrades share more gene families with Arthropoda than with Nematoda. In this network, derived from the OrthoFinder clustering at inflation value 1.5, nodes represent species (0: *Anopheles gambiae*, 1: *Apis mellifera*, 2: *Acyrthosiphon pisum*, 3: *Ascaris suum*, 4: *Brugia malayi*, 5: *Bursaphelenchus xylophilus*, 6: *Caenorhabditis elegans*, 7: *Cimex lectularius*, 8: *Capitella teleta*, 9: *Dendroctonus ponderosae*, 10: *Daphnia pulex*, 11: *Hypsibius dujardini*, 12: *Ixodes scapularis*, 13: *Meloidogyne hapla*, 14: *Nasonia vitripennis*, 15: *Octopus bimaculoides*, 16: *Priapulus caudatus*, 17: *Pediculus humanus*, 18: *Plectus murrayi*, 19: *Pristionchus pacificus*, 20: *Plutella xylostella*, 37: *Ramazzottius varieornatus*, 22: *Solenopsis invicta*, 23: *Strigamia maritima*, 24: *Tribolium castaneum*, 25: *Trichuris muris*, 26: *Trichinella spiralis*, 27: *Tetranychus urticae*, 38: *Drosophila melanogaster*). The thickness of the edge connecting two nodes is weighted by the count of shared occurrences of both nodes in OrthoFinder-clusters. Links involving *H. dujardini* (red) and *R. varieornatus* (orange) are highlighted in colour. The inset box on the lower right shows the average weight of edges between each phylum and both Tardigrades, normalised by the maximum weight (*i.e*. count of co-occurrences of Tardigrades and the annelid *C. teleta*)" **C** Gene family birth synapomorphies at key nodes in Ecdysozoa under two hypotheses: ((Nematoda,Tardigrada),(Arthropoda)) *versus* ((Nematoda),(Tardigrada,Arthropoda)). Each graph shows the number of gene families at the specified node inferred using Dollo parsimony from OrthoFinder clustering at inflation value 1.5. Gene families are grouped by the proportion of taxa above that node that contain a member. Note that to be included as a synapomorphy of the node, the gene family must contain at least one representative each from species at either side of the first child node above the analysed node, and thus there are no synapomorphies with <0.3 proportional proteome coverage in Nematoda and <0.2 in Arthropoda, and all synapomorphies of Tardigrada have 1.0 representation. **D** Gene family birth synapomorphies for Tardigrada+Arthropoda (grey) and Tardigrada+Nematoda (yellow) for OrthoFinder clusterings performed at different MCL inflation parameters.

In *H. dujardini*, a reduced HOX gene complement (six genes in five orthology groups) has been reported [70], and we confirmed this reduction using our improved genome (Figure 5A). The same, reduced complement was also found in the genome of *R. varieornatus* [22], and the greater contiguity of the *R. varieornatus* genome shows that five of the six HOX loci are on one large scaffold, distributed over 2.7 Mb, with 885 non-HOX genes separating them. The *H. dujardini* loci were unlinked in our assembly, except for the two AbdB-like loci, and lack of gene level synteny precludes simple linkage of these scaffolds based on the *R. varieornatus* genome. The order of the HOX genes on the *R. varieornatus* scaffolds is not collinear with other, unfragmented clusters, as *ftz* and the pair of *AbdB* genes are inverted, and *dfd* is present on a second scaffold (and not found between *hox3* and *ftz* as would be expected). The absences of *pb, scr* and *Ubx/AbdA* in both tardigrade species is reminiscent of the situation in Nematoda, where these loci are also absent [71–73].

HOX gene evolution in Nematoda has been dynamic (Figure 5A). No Nematode HOX2 or HOX5 orthology group genes were identified, and only a few species had a single HOX6-8 orthologue. Duplication of the HOX9/AbdB locus was common, generating, for instance the *egl-5, php-3* and *nob-1* loci in *Caenorhabditis* species. The maximum number of HOX loci in a nematode was seven, deriving from six of the orthology groups. Loss of HOX3 happened twice (in *Syphacia muris* and in the last common ancestor of Tylenchomorpha and Rhabditomorpha). The independent loss in *S. muris* was confirmed in two related pinworms, *Enterobius vermicularis* and *Syphacia oblevata*. The pattern of presence and absence of the *Antp* like HOX6-8 locus is more complex, requiring six losses (in the basally-arising enoplean *Enoplis brevis*, the chromadorean *Plectus sambesii*, the pinworm *Syphacia muris*, the ancestor of Tylenchomorpha, the diplogasteromorph *Pristionchus pacificus*, and the ancestor of *Caenorhabditis)*. We affirmed loss in the pinworms by screening the genomes of *E. vermicularis* and *S. oblevata* as above, and no Antp-like locus is present in any of the over 20 genomes available for *Caenorhabditis*. A PCR survey for HOX loci and screening of a *de novo* assembled transcriptome from the nematomorph *Paragordius varius* identified six putative loci from five HOX groups. The presence of a putative HOX2/pb-like gene suggests that loss of HOX2 may be independent in Tardigrada and Nematoda.

Gene family birth can be used as another rare genomic marker. We analyzed the whole proteomes of ecdysozoan taxa for gene family births that supported either the Tardigrada+Nematoda model or the Panarthropoda (Tardigrada+Arthropoda) model. We mapped gene family presence and absence across the two contrasting phylogenies using kinfin v0.8.2 [60] using different inflation parameters in the MCL step in OrthoFinder (Supplementary Data S6:Orthofiner.clustering.tar.gz). Using the default inflation value (of 1.5) the two Tardigrades shared more gene families with Arthropoda than they did with Nematoda (Figure 5B). The number of gene family births synapomorphic for Arthropoda and Nematoda were identical under both phylogenies, as was expected (Table 3; Figure 5C; See Supplementary Data S7: KinFin_output.tar.gz). Many synapomorphic families had variable presence in the daughter taxa of the last common ancestors of Arthropoda and Nematoda, likely because of stochastic gene loss or lack of prediction. However, especially in Nematoda, most synapomorphic families were present in a majority of species (Figure 5C).

**Table 3.** Gene family births that support different relationships of Tardigrada.

| Family_id | Number<br>of<br>proteins | Proportion of proteomes represented |  |  |  | Domain annotations* |
| --- | --- | --- | --- | --- | --- | --- |
|  |  | all | Nematoda<br>(n=9) | Arthropoda<br>(n=15) | Tardigrada<br>(n=2) |  |
| Synapomorphies with membership ≥ 0.7 under the Panarthropoda (Tardigrada + Arthropoda) hypothesis |  |  |  |  |  |  |
| OG0000436 | 104 | 1.00 | 0.00 | 1.00 | 1.00 | Serine proteases, trypsin domain (IPR001254) |
| OG0001236 | 54 | 1.00 | 0.00 | 1.00 | 1.00 | Major facilitator superfamily associated domain (IPR024989) |
| OG0002592 | 36 | 1.00 | 0.00 | 1.00 | 1.00 | Spaetzle (IPR032104) |
| OG0006538 | 19 | 1.00 | 0.00 | 1.00 | 1.00 | Leucine-rich repeat (IPR001611) |
| OG0006541 | 19 | 1.00 | 0.00 | 1.00 | 1.00 | None |
| OG0006869 | 17 | 1.00 | 0.00 | 1.00 | 1.00 | Thioredoxin domain (IPR013766) |
| OG0005117 | 27 | 0.88 | 0.00 | 0.93 | 0.50 | BTB/POZ domain (IPR000210) |
| OG0005941 | 22 | 0.77 | 0.00 | 0.73 | 1.00 | None |
| OG0006662 | 18 | 0.82 | 0.00 | 0.80 | 1.00 | None |
| OG0006889 | 17 | 0.71 | 0.00 | 0.73 | 0.50 | None |
| OG0006940 | 17 | 0.82 | 0.00 | 0.87 | 0.50 | EGF-like domain (IPR000742), Laminin G domain (IPR001791) |
| OG0006941 | 17 | 0.71 | 0.00 | 0.67 | 1.00 | EF-hand domain (IPR002048) |
| OG0006951 | 17 | 0.71 | 0.00 | 0.67 | 1.00 | Adipokinetic hormone (IPR010475) |
| OG0007141 | 16 | 0.82 | 0.00 | 0.80 | 1.00 | None |
| OG0007285 | 15 | 0.71 | 0.00 | 0.67 | 1.00 | GPCR, family 2, secretin-like (IPR000832) |
| OG0007290 | 15 | 0.82 | 0.00 | 0.80 | 1.00 | Allatostatin (IPR010276) |
| OG0007298 | 15 | 0.88 | 0.00 | 0.87 | 1.00 | None |
| OG0007328 | 15 | 0.71 | 0.00 | 0.67 | 1.00 | Sulfakinin (IPR013259) |
| OG0007463 | 14 | 0.77 | 0.00 | 0.73 | 1.00 | Peptidase S1A, nudel (IPR015420), Serine proteases, trypsin domain (IPR001254), Low-density lipoprotein (LDL) receptor class A repeat (IPR002172) |
| OG0007689 | 13 | 0.71 | 0.00 | 0.67 | 1.00 | Marvel domain (IPR008253) |
| Synapomorphies with membership ≥ 0.7 under the Tardigrada + Nematoda hypothesis |  |  |  |  |  |  |
| OG0005423 | 26 | 0.82 | 0.89 | 0.00 | 0.50 | Amidinotransferase (PF02274) |
| OG0006414 | 20 | 0.82 | 0.78 | 0.00 | 1.00 | Proteolipid membrane potential modulator (IPR000612) |
| OG0007199 | 16 | 0.91 | 1.00 | 0.00 | 0.50 | Zona pellucida domain (IPR001507) |
| OG0007812 | 13 | 0.82 | 0.78 | 0.00 | 1.00 | None |
| OG0008368 | 11 | 0.82 | 0.78 | 0.00 | 1.00 | RUN domain (IPR004012) |

\* Domain annotations are reported where proteins from more than one third of the proteomes in the family had that annotation. IPR = InterPro domain identifier; PF = PFam identifier.

At inflation value 1.5, we found six gene families present that had members in both tardigrades and all 14 arthropods under Panarthropoda, but no gene families were found in both tardigrades and all 9 nematodes under the Tardigrada-Nematoda hypothesis (Supplemental Table 11). Allowing for stochastic absence, we inferred 154 families to be synapomorphic for Tardigrada+Arthropoda under the Panarthropoda hypothesis, and 99 for Tardigrada-Nematoda under the Tardigrada+Nematoda hypothesis (Figure 5D). More of the Tardigrada+Arthropoda synapomorphies had high species representation than did the Tardigrada-Nematoda synapomorphies. This pattern was also observed in analyses using different inflation values and in analyses including the transcriptome from the nematomorph *Paragordius varius*.

At inflation value 1.5, we found six gene families present that had members in both tardigrades and all 14 arthropods under Panarthropoda, but no gene families were found in both tardigrades and all 9 nematodes under the Tardigrada+-Nematoda hypothesis (Table 3). Allowing for stochastic absence, we inferred 154 families to be synapomorphic for Tardigrada+Arthropoda under the Panarthropoda hypothesis, and 99 for Tardigrada+-Nematoda under the Tardigrada+-Nematoda hypothesis (Figure 5D). More of the Tardigrada+Arthropoda synapomorphies had high species representation than did the Tardigrada+- Nematoda synapomorphies. This pattern was also observed in analyses using different inflati on values and in analyses including the transcriptome from the nematomorph *Paragordius varius* (Table 3).

We explored the biological implications of these putative synapomorphies by examining the functional annotations of each protein family that had ≥70% of the ingroup species represented Table 3. Under Tardigrada+Nematoda, five families were explored. Four of these had domain matches (proteolipid membrane potential modulator, zona pellucida, RUN and amidinotransferase domains), and one did not contain any protein with an identifiable domain.

Under Panarthropoda, twenty families had ≥70% of the ingroup taxa represented, and six were universally present. These included important components of developmental and immune pathways, neuromodulators and others. Of particular note, two families, one universal, were annotated as serine endopeptidases, one including *D. melanogaster* Nudel, and another universal family included Spätzle orthologs. Spätzle is a cysteine-knot, cytokine-like ligand involved in dorso-ventral patterning, and is the target of a serine protease activation cascade initiated by Nudel protease. In *D. melanogaster*, Nudel is a maternally supplied signal that acts with other genes to activate Easter, a protease that cleaves Spätzle. Spätzle interacts with the Toll receptor pathway. The identification of several members of a single regulatory cascade as potential gene family births suggests that the pathway may have been established in the Tardigrade-Arthropod last common ancestor. Other Panarthropoda-synapomorphic families were annotated with *Drosophila* adipokinetic hormone, neuromodulatory allatostatin-A and drosulfakinin, leucine-rich repeat, thioredoxin, major facilitator superfamily associated, and “domain of unknown function” DUF4728 domains. Eight of the fourteen Panarthropoda synapomorphic families had no informative domain annotations. One family included orthologues of *Drosophila* eyes-shut, a component of the, a component of the apical extracellular matrix of the ommatidial retina. Other Panarthropoda-synapomorphic families were annotated with adipokinetic hormone, neuromodulatory allatostatin-A, drosulfakinin, leucine-rich repeat, thioredoxin, major facilitator superfamily associated, and “domain of unknown function” DUF4728 domains. However, nine of the twenty Panarthropoda synapomorphic families had no informative domain annotations.

## DISCUSSION

### A ROBUST ESTIMATE OF THE *HYPSIBIUS DUJARDINI* GENOME

We have sequenced and assembled a high quality genome for the tardigrade *H. dujardini*, utilizing new data, including single-molecule sequencing long-read, and heterozygosity aware assembly methods. Comparison of genomic metrics with previous assemblies for this species showed that our assembly is much more contiguous than has been achieved previously, and retains minimal uncollapsed heterozygous regions. Furthermore, the lack of suspiciously high coverage scaffolds and the low duplication rate of CEGMA genes implies a low rate of scaffold over-assembly. The span of this new assembly is much closer to independent estimates of the size of the *H. dujardini* genome (75 - 100 Mb) using densitometry and staining.

The *H. dujardini* genome is thus nearly twice the size of that of the related tardigrade *R. varieornatus*. We compared the two genomes to identify differences that would account for the larger genome in *H. dujardini*. While we predict *H. dujardini* to have ~6,000 more protein coding genes than *R. varieornatus*, these account for only ~23 Mb of the additional span, and are not obviously simple duplicates of genes in *R. varieornatus*. Analyses of the gene contents of the two species showed that while *H. dujardini* had more species-specific genes, it also had greater numbers of loci in species-specific gene family expansions than *R. varieornatus*, and had lost fewer genes whose origins could be traced to a deeper ancestor. *H. dujardini* genes had, on average, the same structure (~6 exons per gene) as did *R. varieornatus*, however introns in *H. dujardini* orthologous genes were on average twice the length of those in *R. varieornatus* (255 bases versus 158 bases). Finally, the *H. dujardini* genome was more repeat rich (26% compared to only 18% in *R. varieornatus)*. These data argue against simple whole genome duplication in *H. dujardini*. The genome of *H. dujardini* is larger because of expansion of non-coding DNA, including repeats and introns, and acquisition and retention of more new genes and gene duplications than *R. varieornatus*. The disparity in retention of genes with orthologues outside the Tardigrada, where *R. varieornatus* has lost more genes than has *H. dujardini*, suggests that *R. varieornatus* may have been selected for genome size reduction, and that the ancestral tardigrade (or hypsibiid) genome is more likely to have been ~100 Mb than 54 Mb. We await additional tardigrade genomes with interest.

While we identified linkage between genes in the two tardigrades, local synteny was relatively rare. In this these genomes resemble those of the genus *Caenorhabditis*, where extensive, rapid, within-chromosome rearrangement has served to break close synteny relationships while, in general, maintaining linkage [74]. We have found that a high proportion of the orthologs of genes on the same scaffold on *H. dujardini* are on the same scaffold of *R. varieornatus* as well, which supports intrachromosomal rearrangements. The absence of chromosomal level assemblies for either tardigrade (and lack of any genetic map information) precludes definitive testing of this hypothesis.

### HORIZONTAL GENE TRANSFER IN TARDIGRADES: *H. DUJARDINI* HAS A NORMAL METAZOAN GENOME

Boothby *et al.* made the surprising assertion that 17.5% of *H. dujardini* genes originated through HGT from a wide range of bacterial, fungal and protozoan donors. Subsequently, several groups including our teams proved that this finding was the result of contamination of their tardigrade samples with cobionts, and less-than-rigorous screening of HGT candidates. We found that the use of uncurated gene-finding approaches also yielded elevated HGT proportion estimates in many other nematode and arthropod genomes, even where contamination is unlikely to be an issue. It is thus essential to follow up initial candidate sets with detailed validation steps. We screened our new *H. dujardini* assembly for evidence of HGT, identifying a maximum of 3.7% of the protein coding genes as potential candidates. After careful assessment using phylogenetic analysis and expression evidence, we identified a maximum of 2.3% and a likely high-confidence set of only 0.6% of *H. dujardini* genes that originated through HGT. HGT was also much reduced (1.6%) in the high-quality *R. varieornatus* genome. These proportions are congruent with similar analyses of *C. elegans* and *D. melanogaster*. Curation of the genome assemblies and gene models may decrease the proportion further. Tardigrades do not have elevated levels of HGT.

While the tardigrades do not have spectacularly elevated levels of possible HGT in their genomes, some identified HGT events are of importance in anhydrobiosis. All *H. dujardini* catalase loci were of bacterial origin, as described for *R. varieornatus*. While trehalose phosphatase synthase was absent from *H. dujardini*, *R. varieornatus* has a TPS that likely was acquired by HGT. As *M. tardigradum* does not have a TPS homolog, while other ecdysozoan taxa do, this suggests that TPS may have been lost in the common ancestor of eutardigrada and regained in *R. varieornatus* by HGT after divergence from *H. dujardini*.

### CONTRASTING MODES OF ANHYDROBIOSIS IN TARDIGRADES

Protection related genes found in *R. varieornatus* were highly conserved in *H. dujardini*, of which CAHS and SAHS both had high copy numbers. However, we did not find a Dsup ortholog in *H. dujardini*. In addition, *H. dujardini* has similar gene losses in pathways that produce ROS and cellular stress signaling pathways found in *R. varieornatus*, which suggest that the gene losses have occurred before the divergence of the two species. The loss in the major genes of signaling pathways would cause disconnection; various stress inductions may not be relayed to downstream factors, such as cell cycle regulation, transcription and replication inhibition, and apoptosis. In contradiction, various cellular protection and repair pathways are highly conserved, therefore cellular signaling may be inhibited but cellular molecules are protected, and if damaged, repaired.

On the other hand, various gene families have undergone duplication in both tardigrades and linage specifically. In particular, SOD was duplicated in both tardigrades, along with a calcium activated potassium channel, which has been implied to contribute to cellular signaling during anhydrobiosis. Furthermore, we have found that a high number of gene families have been expanded in *H. dujardini* compared to *R. varieornatus*. This may be related to the genome size reduction in *R. varieornatus*, which multiple-copy genes may be selected for loss.

Previous studies have implied that the anhydrobiosis response in the two tardigrades may differ; *H. dujardini* has an induced transcriptomic response where *R. varieornatus* does not. This shows consistency with the phenotype at transfer into anhydrobiosis. *H. dujardini* requires 24 hours at 100% RH prior to desiccation and takes at least 24 hours for a successful desiccation, and *R. varieornatus* can enter anhydrobiosis within 30 minutes at when exposed to 37% RH. It has been implied that genes required for a successful anhydrobiosis are upregulated during the 48 hours of anhydrobiosis induction in *H. dujardini*. Therefore, we have not only sequenced the transcriptome of *R. varieornatus* desiccated in the normal protocol, but also with a slowly dried sample. In a natural tardigrade living environment, desiccation does not progress within a 30-minute time frame, therefore a slowly dried *R. varieornatus* may have induced expression in genes for a more successful anhydrobiosis.

We found that *H. dujardini* has more differentially expressed genes than *R. varieornatus*, which again was supported with a significant increase in distribution of fold change in the whole transcriptome. We have found that a variety of calcium related transporters, receptors to be differentially expressed. Kondo *et al.* suggested that cellular signaling using calmodulin and calcium may be required for anhydrobiosis. Calcium related cellular signaling is used in a variety of cellular signaling, especially in cAMP signaling pathways. It is still unclear if how this calcium induced signaling pathway is related to anhydrobiosis, however the decrease in water molecules would cause an increase in cellular calcium concentration, inducing a signaling cascade. Furthermore, as we anticipated, *R. varieornatus* had higher DEG count when desiccated at a slow pace, which contained CAHS and SAHS genes, and anti-oxidant related genes. Although most of these genes are highly expressed (>100TPM) in the active state, the expression induction of these genes may enable higher recovery ratio.

Unexpectedly, we identified 3 copies of trehalase to be DE of which 2 copies have over 200 TPM in the “tun” state. However, trehalose synthesis pathway via trehalose phosphatase synthase (TPS) has been lost solely in *H. dujardini*. Several studies have determined *M. tardigradum* has also lost the ability to synthesize trehalose, further supported by undetectable levels of trehalose [31]. Trehalose has been known for its protective role of cellular molecules [35, 36, 75, 76], however it has been hypothesized that it is not required for tardigrade anhydrobiosis. Trehalose degradation would not be required if there are no trehalose, therefore there may be a complementary pathway for trehalose synthesis.

Although the two species have obtained different mechanisms for anhydrobiosis induction, we have found several gene families that have increased expression during anhydrobiosis in both *H. dujardini* and slow-dried *R. varieornatus*, such as CAHS and SAHS, predictable candidates of expression induction. We have found calcium related receptors, which are coupled with calcium and cAMP/GMP, as DE. The mechanism of desiccation sensing still remains to be clarified, however the decrease in cellular water molecules would cause an increase in cellular metabolites and metallic ions, *i.e*. calcium, which would be an ideal target for desiccation detection. Furthermore, we have found aquaporin-10 to be highly expressed in *R. varieornatus* and DE in *H. dujardini*. It has been previously reported that *M. tardigradum* has 10 copies of aquaporins [77], which *H. dujardini* has 11, and *R. varieornatus* 10. Aquaporins contribute to transportation of water molecules into cells, which anhydrobiosis induction would be related to [78].

### THE POSITION OF TARDIGRADES IN THE METAZOA

Our phylogenomic analyses found Tardigrada, represented by *H. dujardini* and *R. varieornatus* genomes as well as transcriptomic data from *Milnesium tardigradum* and *Echiniscus testudo*, to be sisters to Nematoda, not Arthropoda. This finding was robust to inclusion of additional phyla, such as Onychophora and Nematomorpha, and to filtering the alignment data to exclude categories of poorly represented or rapidly evolving loci. This finding is both surprising, and not new. Many previous molecular analyses have found Tardigrada to group with Nematoda, whether using single genes or ever larger gene sets derived from transcriptome and genome studies [1–3]. This phenomenon has previously been attributed to long branch attraction in suboptimal datasets, with elevated substitutional rates or biased compositions in Nematoda and Tardigrada mutually and robustly driving Bayesian and Maximum Likelihood algorithms to support the wrong tree. Strikingly, in our analyses including taxa for which transcriptome data are available, we found Onychophora to lie outside a ((Nematoda, Nematomorpha, Tardigrada), Arthropoda) clade. This finding, while present in some other analyses (e.g component phylogenies summarised in [2]), conflicts with accepted systematic and many molecular analyses. We note that Onychophora was only represented by transcriptome datasets, and that there is accordingly an elevated proportion of missing data in the alignment for this phylum.

That a tree linking Tardigrada with Nematoda is “wrong” is a prior supported by developmental and anatomical data: tardigrades are segmented, have appendages, and have a central and peripheral nervous system anatomy that can be homologised with those of Onychophora and Arthropoda [79, 80]. In contrast, nematodes are unsegmented, have no lateral appendages and have a simple nervous system. The myoepithelial triradiate pharynx, found in Nematoda, Nematomorpha, and Tardigrada, is one possible morphological link, but Nielsen has argued persuasively that the structures of this organ in nematodes and tardigrades (and other taxa) are distinct and thus non-homologous [5].

*H. dujardini* has a reduced complement of HOX loci, as does *R. varieornatus*. Some of the HOX loci missing in the Tardigrada are the same as those lost in Nematoda. Whether these absences are a synapomorphy for a Nematode-Tardigrade clade, or simply a product of homoplasious evolution remains unclear. It may be that miniaturisation of Nematoda and Tardigrada during adaptation to life in interstitial habitats facilitated the loss of specific HOX loci involved in post-cephalic patterning, and that both nematodes and tardigrades can be thought to have evolved by reductive evolution from a more fully featured ancestor. While tardigrades retain obvious segmentation, nematodes do not, with the possible exception of repetitive cell lineages along the anterior-posterior axis during development [81]. We note that until additional species were analyzed, the pattern observed in *C. elegans* was assumed to be the ground pattern for all Nematoda. More distantly-related Tardigrada may have different HOX gene complements than these hypsibiids: and a pattern of staged loss similar to that in Nematoda [71–73] may be found. It may be intrinsically easier to lose some HOX loci than others.

Assessment of gene family birth as rare genomic changes lent support to a Tardigrada+Arthropoda clade, but the support was not strong. There were more synapomorphic gene family births when a Tardigrada+Arthropoda (Panarthropoda) clade was assumed than when a Tardigrada+Nematoda clade was assumed. However, analyses under the assumption of Tardigrada+Nematoda identified synapomorphic gene family births at only 50% of the level found when Panarthropoda was assumed. We note that recognition of gene families may be compromised by the same “long branch attraction” issues that plague phylogenetic analyses, and also that any taxon where gene loss is common (such as has been proposed for Nematoda as a result of its simplified body plan may score poorly in gene family membership metrics. The short branch lengths that separate basal nodes in the analysis of the panarthropodan-nematode part of the phylogeny of Ecdysozoa may make robust resolution very difficult. Thus our analyses of rare genomic changes lent some support to the Panarthropoda hypothesis, as did analysis of miRNA gene birth [2], but analysis of HOX loci may conflict with this. For the families that did support Panarthropoda, it was striking that many of these deeply conserved loci have escaped experimental, genetic annotation.

We explored the biological implications of these synapomorphies by examining the functional annotations of each protein family (Supplementary Table S10, Supplemental Data S7). The six loci identified as universally retained gene family births in Panarthropoda included *spaetzle*, a cysteine-knot, cytokine-like family that is known to interact with the Toll receptor pathway in *D. melanogaster*, and is (in that species) involved in dorso-ventral patterning as well as immune response. Other clusters were functionally annotated as having serine-type endopeptidase activity (this is not nudel, but nudel is also a Serine proteases, trypsin domain) or harboring a thioredoxin domain and thus being involved in cell redox homeostasis. However the remainder of the clusters had no informative annotation other than the presence of “domains of unknown function” (DUFs). the arthropod specific “domain of unknown function” DUF4728, a major facilitator superfamily associated domain and a Leucine-rich repeat domain. It is striking that such deeply conserved loci should have escaped functional, genetic or biochemical annotation.

By lowering the threshold of proteome coverage to 70% when declaring synapomorphies, synapomorphic clusters can found under the Triradiata-hypothesis (5 of which 3 contain both Tardigrades), but are fewer than under the Panarthropoda hypothesis (14 of which 11 contain both Tardigrades). 70%-Synapomorphies under Triradiata contains clusters representing Proteolipid membrane potential modulator domains (both Tardigrades), Zona pellucida domains (both Tardigrades), RUN domains (both Tardigrades), Amidinotransferase domains, while one cluster did not contain any protein with an identifiable domain. Of the 14 70%-Synapomorphies under Panarthropoda, 8 did either not contain any proteins with identifiable domains or contained uncharacterized *Drosophila* orthologues. One cluster was formed by *Drosophila* eyes shut orthologues (missing in *R. varieornatus)*, essential for the formation of matrix-filled interrhabdomeral space, maybe important contribution to insect eye evolution). One cluster contained *Drosophila* Adipokinetic hormone orthologues (missing in *R. varieornatus*), which is associated with marked increase in hemolymph carbohydrate. One cluster contained *Drosophila* Allatostatin-A orthologues, which may act as a neurotransmitter or neuromodulator. One Drosulfakinins orthologue, which plays diverse biological roles including regulating gut muscle contraction in adults but not in larvae. And Serine protease nudel orthologues, which is also an component of the extracellular signaling pathway that establishes the dorsal-ventral pathway of the embryo, and acts together with gd (gastrulation-defective) and snk (snake) to process easter into active easter (drosophila easter and snake are recovered in the fourth biggest cluster, containing representatives of all proteomes, gd is in cluster containing 2 Nematodes and 11 Arthropods), which subsequently defines cell identities along the dorso-ventral continuum by activating the spz ligand for the toll receptor in the ventral region of the *Drosophila* embryo.

We regard the question of tardigrade relationships to be open. While a clade of Tardigrada, Onychophora, Arthropoda, Nematoda and Nematomorpha was supported, the branching order within this group remains contentious, and in particular the positions on Tardigrada and Onychophora are poorly supported and/or variable in our and others’ analyses. Full genome sequences of representatives of Onychophora, Heterotardigrada (the sister group to the Eutardigrada including Hypsibiidae), Nematomorpha and enoplian (basal) Nematoda are required. Resolution of the conflicts between morphological and molecular data will be informative - either of the ground state of a nematode-tardigrade ancestor, or of the processes that drive homoplasy in “rare” genomic changes and robust discovery of non-biological trees in phylogenomic studies.

## METHODS

### TARDIGRADE CULTURE AND SAMPLING

The tardigrade *Hypsibius dujardini* Z151 was purchased from Sciento (Manchester, UK). *H. dujardini* ZI5I and *Ramazzottius varieornatus* strain YOKOZUNA-1 were cultured as previously described [24, 42]. Briefly, tardigrades were fed *Chlorella vulgaris* (Chlorella Industry) on 2% Bacto Agar (Difco) plates prepared with Volvic water, incubated at 18°C for *H. dujardini* and 22°C for *R. varieornatus* under constant dark conditions. Culture plates were renewed every 7~8 days. Anhydrobiotic adult samples were isolated on 30μΜ filters (MILLIPORE), and placed in a chamber maintained at 85% relative humidity (RH) for 48hr for *H. dujardini*, and 30% RH for 24 hr and additional 24 hr at 0% RH for slow-dried *R. varieornatus*, and 0% RH for 30 minutes on a 4 cm × 4 cm Kim-towel with 300μL of distilled water, and additional 2 hours without the towel for fast-dried *R. varieornatus*. Successful anhydrobiosis was assumed when >90% of the samples prepared in the same chamber recovered after rehydration.

### SEQUENCING

Genomic DNA for long read sequencing was extracted using MagAttract HMW DNA Kit (Qiagen) from approximately 900,000 *H. dujardini*. DNA was purified twice with AMPure XP beads (Beckman Coulter). A 20 kb PacBio library was prepared following the manual “20 kb Template Preparation Using BluePippin Size-Selection System (15 kb Size Cutoff)” (PacBio SampleNet - Shared Protocol) using SMARTBell Template Prep Kit 1.0 (Pacific Biosciences), and was sequenced using 8 SMRT Cells Pac V3 with DNA Sequencing Reagent 4.0 on a PacBio RSII System (Pacific Biosciences) at Takara Bio Inc. Briefly, purified DNA was sheared, targeting 20 kb fragments, using a g-TUBE (Covaris). Following end-repair and ligation of SMRTbell adapters, the library was size-selected using BluePippin (Sage Science) with a size cut-off of 10 kb. The size distribution of the library was assayed on TapeStation 2200 (Agilent) and quantified using the Quant-iT dsDNA BR Assay Kit (Invitrogen). MiSeq reads from a single *H. dujardini* individual (DRR055040) are from our previous report [21].

For gene prediction RNA-Seq, 30 individuals were collected from each of the following conditions in three replicates: active and dried adults (slow dried for *R. varieornatus)*, eggs (1, 2, 3, 4, 5, 6 and 7 days after laying) and juveniles (1, 2, 3, 4, 5, 6 and 7 days after hatching). Due to sample preparations, *R. varieornatus* juveniles were sampled until juvenile 1st day. In addition for gene expression analysis, we sampled 2~3 individuals for fast-dried *R. varieornatus*. All RNA-Seq analysis was conducted with 3 replicates. Specimens were thoroughly washed with Milli-Q water on a sterile nylon mesh (Millipore), immediately lysed in TRIzol reagent (Life Technologies) using three cycles of immersion in liquid nitrogen followed by 37°C incubation. Total RNA was extracted using the Direct-zol RNA kit (Zymo Research) following the manufacturer’s instructions, and RNA quality was checked using the High Sensitivity RNA ScreenTape on a TapeStation (Agilent Technologies). For library preparation, mRNA was amplified using the SMARTer Ultra Low Input RNA Kit for Sequencing v.4 (Clontech), and Illumina libraries were prepared using the KAPA HyperPlus Kit (KAPA Biosystems). Purified libraries were quantified using a Qubit Fluorometer (Life Technologies), and the size distribution was checked using the TapeStation D1000 ScreenTape (Agilent Technologies). Libraries size selected above 200 bp by manually excision from agarose were purified with a NucleoSpin Gel and PCR Clean-up kit (Clontech). The samples were then sequenced on the Illumina NextSeq 500 in High Output Mode with a 75-cycle kit (Illumina) as single end reads, with 48 multiplexed samples per run. Adapter sequences were removed, and sequences were demultiplexed using the bcl2fastq v.2 software (Illumina). For active and dried adults, RNA-Seq was also conducted starting from approximately 10,000 individuals, similarly washed but RNA extraction with TRIzol reagent (Life Technologies) followed by RNeasy Plus Mini Kit (Qiagen) purification. Library preparation and sequencing was conducted at Beijing Genomics Institute.

For miRNA-Seq, 5,000 individuals were homogenized using Biomasher II (Funakoshi), and TRIzol (Invitrogen) was used for RNA extraction, and purified by isopropanol precipitation. Size selection of fragments of 18-30 nt using electrophoresis, preparation of the sequencing library for Illumina HiSeq 2000 and subsequent (single end) sequencing was carried out by Beijing Genomics Institute.

All sequenced data were validated for quality by FastQC [82].

### GENOME ASSEMBLY

The MiSeq reads from the WGA were merged with Usearch [83] and both merged and unmerged pairs were assembled with SPAdes [84] as single-end. The SPAdes assembly was checked for contamination with BLAST+ BLASTN [85] against the nr [86] database and no observable contamination was found with blobtools [87]. The UNC Illumina libraries were mapped to the SPAdes assembly with Bowtie2 [88] and read pairs were retained if at least one of them mapped to the assembly. These reads were then assembled, scaffolded and gap closed with Platanus [44]. The Platanus assembly was further scaffolded and gap closed using the PacBio data with PBJelly [89].

Falcon [43] assembly was performed on the DNAnexus platform. Using this Falcon assembly, Platanus assembly was extended using SSPACE-LongReads [46], and gap-filled with PBJelly [89] with default parameters. Single individual MiSeq reads were mapped to the assembly, and all contigs with coverage < 1, length < 1000, or those corresponding to the mitochondrial genome were removed. At this stage, one CEGMA gene became unrecognized by CEGMA [47] probably due to multiple PBJelly runs, and therefore the contig harboring that missing CEGMA gene was corrected by Pilon [90] using the single individual MiSeq reads.

### GENE FINDING

Prior to the annotation, mRNA-Seq data (Development, Active-tun 10k animals) were mapped to the genome assembly with TopHat [88, 91] without any options. Using the mapped data from TopHat, BRAKER [48] was used with default settings to construct a GeneMark-ES [92] model and an Augustus [61] gene model, which are used for *ab initio* prediction of genes. The getAnnotFasta.pl script from Augustus was used to extract coding sequences from the GFF3 file. Similarly, to construct a modified version of the *R. varieornatus* genomes annotation, we used the development and anhydrobiosis (Supplementary Table S1BC) RNA-Seq data for BRAKER annotation. We found that several genes were mis-annotated (MAHS in both species, a CAHS ortholog in *R. varieornatus*), mainly fused with a consecutive gene, therefore manual curation were conducted. tRNA and rRNA genes were predicted with tRNAscan-SE [51] and RNAmmer [50], respectively.

The RNA-Seq data used to predict the gene models were mapped with BWA MEM [93] against the predicted CDS sequences, the genome, and a Trinity [49] assembled transcriptome. We also mapped the RNA-Seq data used for gene expression analysis (single individual *H. dujardini* and fast/slow dry of *R. varieornatus*) of the active state and tun state. After SAM to BAM conversion and sorting with SAMtools view and sort [94], we used QualiMap [95] for mapping quality check.

To annotate the predicted gene models with genes from published databases, we ran a similarity search using BLAST v2.2.22 BLASTP [64] against Swiss-Prot, TrEMBL [57], and HMMER hmmsearch [96] against Pfam-A [97] and Dfam [56], KAAS analysis for KEGG ortholog mapping [98], and InterProScan [99] for domain annotation. We used RepeatScout [100] and RepeatMasker [101] for de novo repeat identification. Furthermore, in order to compare the gene models of those of *R. varieornatus*, we also ran BLAST v2.2.22 BLASTP searches against the updated *R. varieornatus* proteome, and TBLASTN search against the *R. varieornatus* genome. Additionally, we determined tight link orthologs between *H. dujardini* and *R. varieornatus* orthologs using BLAST v2.2.22 BLASTP, and extracted bidirectional best hits with in-house Perl scripts.

For miRNA prediction we used miRDeep [52] to predict mature miRNA within the genome, using the mature miRNA sequences in miRBase [53]. The predicted mature miRNA sequences were then searched against miRBase with ssearch36 [102] for annotation by retaining hits with identity > 70% and a complete match of bases 1-7, 2-8 or 3-9.

### HORIZONTAL GENE TRANSFER ANALYSES

HGT genes were determined on the HGT index [62]. Both Swiss-Prot and TrEMBL data of the UniProt division database was downloaded [57], and sequences with “Complete Proteome” in the Keyword were extracted for following analysis. Following the method of Boschetti *et al.*, an Arthropoda-less and Nematoda-less database was constructed as well as the whole database, for excluding similarity search hits of the same phylum for the query organism. A DIAMOND [63] database was constructed, and the CDS sequences were subjected to DIAMOND BLASTX. Genes with no hits with an E-value below 1e-5 were filtered out. The HGT index (Hu) was calculated by Bm - Bo, the bit score difference between the best non-metazon hit (Bo) and the best metazoan hit (Bm), and genes with Hu ≥ 30 were identified as a HGT candidate. The HGT percentage was calculated by the number of genes with Hu ≥ 30 divided by the number of genes with a 1e-5 hit. The longest transcript for each gene was used in this analysis to exclude splicing variations.

To assess if *ab initio* annotation of genomes has a bias in the calculation of HGT index, we calculated the HGT index for genomes in ENSEMBL-metazoa [54] that had a corresponding Augustus v3.2.2 [61] gene model to run *ab initio* gene prediction for comparison (*Aedes aegypti*, *Apis mellifera*, *Bombus impatiens*, *Caenorhabditis brenneri*, *C. briggsae*, *C. elegans*, *C. japonica*, *C. remanei*, *Culex quinquefasciatus*, *Drosophila ananassae*, *D. erecta*, *D. grimshawi*, *D. melanogaster*, *D. mojavensis*, *D. persimilis*, *D. pseudoobscura*, *D. sechellia*, *D. simulans*, *D. virilis*, *D. willistoni*, *D. yakuba*, *Heliconius melpomene*, *Nasonia vitripennis*, *Rhodnius prolixus*, *Tribolium castaneum*, *Trichinella spiralis*). The genome of *R. varieornatus* [22] were downloaded from their corresponding genome projects, and submitted for analysis. Gene predictions for each organism were conducted using the script autoAugPred.pl of the Augustus package with the corresponding model (Supplementary Table S8). The longest isoform sequence for all genes were extracted for both ENSEMBL and *ab initio* annotations, and HGT index was calculated for each gene in all organisms. Furthermore, in order to assess if using DIAMOND BLASTX biased the calculation, we ran BLAST v2.2.22 BLASTX [64] searches with *H. dujardini*, and calculated the HGT index using the same pipeline.

The blast-score based HGT index provided a first-pass estimate of whether a gene had been horizontally transferred from a non-metazoan species. Phylogenetic trees were constructed for each of the 463 candidates (based on the HGT index) along with their best blast hits as described above (Supplementary Data S3: hgt_trees.tar.gz). Protein sequences for the blast hits were aligned along with the HGT candidate using MAFFT [103]. RAxML [65] was used to build 461 individual trees as 2 of the protein sets had less than 4 sequences and trees could not be built for them. HGT candidates were categorized as prokaryotes, viruses, metazoan, and non-metazoan (i.e., eukaryotes that were non-metazoan, such as fungi) based on the monophyletic clades that they were placed in. Any that could not be classified monophyletically were classified as complex (Supplementary Data S8: Supp_data_S8_463_putative HGTs). OrthoFinder [59] with default BLAST+ BLASTP search settings and an inflation parameter of 1.5 was used to identify orthogroups containing *H. dujardini* genes and *R. varieornatus* protein-coding genes. These orthogroups were used to identify the *R. varieornatus* HGT homologs of *H. dujardini* HGT candidates. HGT candidates were also classified as having high gene expression levels if they had an average gene expression greater than the overall average gene expression level of 1 TPM.

### ANHYDROBIOSIS ANALYSES

To identify the genes responsive to anhydrobiosis, we explored transcriptome (Illumina RNA-Seq) data for both *H. dujardini* and *R. varieornatus*. Individual RNA-Seq data for *H. dujardini* [42] before and during anhydrobiosis were contrasted with new sequence data for *R. varieornatus* similarly treated. We mapped the RNA-Seq reads to the coding sequences of the relevant species with BWA MEM [93] and after summarizing the read count of each gene, we used DESeq2 [104] for differential expression calculation, using false discovery rate (FDR) correction. Genes with a FDR below 0.05, an average expression level (in transcripts per kilobase of model per million mapped fragments; TPM) of over 1, and a fold change over 2 were defined as differentially expressed genes. Gene expression (TPM) was calculated with Kallisto [105], and was parsed with custom Perl scripts. To assess if there were any differences in fold change distributions, we used R to calculate the fold change for each gene ( anhydrobiotic / (active+0.1)), and conducted a ⋃ test using the wilcox.text() function. We mapped the differentially expressed genes to KEGG pathway maps [106] to identify pathways that were likely to be differentially active during anhydrobiosis.

### PROTEIN FAMILY ANALYSES and COMPARATIVE GENOMICS

For comparison with *R. varieornatus*, we first aligned the genomes of *H. dujardini* and *R. varieornatus* with Murasaki, and visualized with gmv [58]. The lower tf-idf anchor filter was set to 500. A syntenic block was seen between scaffold0001 of *H. dujardini* and Scaffold002 of *R. varieornatus*, therefore, we extracted the corresponding regions (*H. dujardini:* scaffold0001 363,334-2,100,664, *R. varieornatus:* Scaffold002 2,186,607-3,858,816), and conducted alignment with Mauve [107]. Furthermore, to validate the tendency whether intra- or inter-chromosomal rearrangements were selected, we determined the number of bidirectional best hit (BBH) orthologs on the same scaffold in both *H. dujardini* and *R. varieornatus*. In addition, we extracted gene pairs that had an identity of more than 90% by ClustalW2 [99], and calculated the identity of first and last exon between pairs. Tardigrade specific protection related genes (CAHS, SAHS, MAHS, RvLEAM, Dsup) were identified by BLASTP against TrEMBL, and were subjected to phylogenetic analysis using Clustalw2 [99] and FastTree [108], and visualized with FigTree [109].

HOX loci were identified using BLAST, and their positions on scaffolds and contigs assessed. To identify HOX loci in other genomes, genome assembly files were downloaded from ENSEMBL Genomes [54] or Wormbase ParaSite [110, 111] and formatted for local search with BLAST+ [85]. Homeodomain alignments were generated using Clustal Omega [112] and phylogenies estimated with RAxML [65].

Protein predictions from genomes of Annelida (1), Nematoda (9), Arthropoda (15), Mollusca (1), Priapulida (1) were clustered together with the proteins of *H. dujardini* and *R. varieornatus* using OrthoFinder [59] at different inflation values (Supplementary Data S6: Orthofinder.clustering.tar.gz). OrthoFinder output (Supplementary Data 7: KinFin_input.tar.gz) was analyzed using KinFin v0.8.2 [60] using the attribute file under two competing phylogenetic hypotheses: either “Panarthropoda”, where Tardigrada and Arthropoda share a LCA or where Tardigrada and Nematoda share a LCA. Enrichment and depletion in clusters containing proteins from Tardigrada and other taxa was tested using a two-sided Mann-Whitney-U test of the count (if at least two taxa had non-zero counts) and results were deemed significant at a p-value threshold of p=0.01.

Protein predictions from genomes of Annelida (1), Nematoda (9), Arthropoda (15), Mollusca (1), Priapulida (1) were retrieved from public databases (see Supplementary Table S7 for proteome sources). Proteomes were screened for isoforms (Supplementary Data S10: proteome_fastas) and longest isoforms were clustered with the proteins of *H. dujardini* and *R. varieornatus* using OrthoFinder 1.1.2 [59] at different inflation values. Proteins from all proteomes were functionally annotated using InterProScan [99].

OrthoFinder output (Orthologous_groups.txt) was analysed using KinFin v0.8.2 [60] under two competing phylogenetic hypotheses: either “Panarthropoda”, where Tardigrada and Arthropoda share a LCA or where Tardigrada and Nematoda share a LCA (see Supplementary Data S7: KinFin_input.tar.gz for input files used in KinFin analysis). Within KinFin, Enrichment and depletion in clusters containing proteins from Tardigrada and other taxa was tested using a two-sided Mann-Whitney-⋃ test of the count (if at least two taxa had non-zero counts) and results were deemed significant at a p-value threshold of p=0.01.

A graph-representation of the OrthoFinder clustering (at Inflation value = 1.5) was generated using the generate_network.py script distributed with KinFin v0.8.2. The nodes in the graph were positioned using the ForceAtlas2 layout algorithm implemented in Gephi v0.9.1 (“Scaling” 10000.0, “Stronger Gravity” = True, “Gravity” = 1.0, “Dissuade hubs” = False, “LinLog mode” = True, “Prevent overlap” = False, “Edge Weight Influence” = 1.0).

Single-copy orthologues between *H. dujardini* and *R. varieornatus* were identified using the orthologous groups defined by OrthoFinder. Using the Ensembl Perl API, gene structure information (gene lengths, exon counts and intron spans per gene) were extracted for these gene pairs. To avoid erroneous gene predictions biasing observed trends, *H. dujardini* genes which were 20% longer or 20% shorter were considered outliers.

### PHYLOGENOMICS

In order to improve phylogenetic coverage, transcriptome data was retrieved for additional tardigrades (2), a priapulid (1), kinorhynchs (2) and onychophorans (3) (see Supplementary Table S15). Assembled transcripts for *Echiniscus testudo, Milnesium tardigradum, Pycnophyes kielensis* and *Halicryptus spinulosus* were downloaded from NCBI Transcriptome Shotgun Assembly (TSA) Database. EST sequences of *Euperipatoides kanangrensis, Peripatopsis sedgwicki* and *Echinoderes horni* were download from NCBI Trace Archive and assembled using CAP3 [113]. Raw RNA-seq reads for *Peripatopsis capensis* were downloaded from NCBI SRA, trimmed using skewer v0.2.2 [114] and assembled with Trinity v2.2.0 [49]. Protein sequences were predicted from all transcriptome data using TransDecoder [115] (retaining a single open reading frame per transcript). Predicted proteins were used in an additional OrthoFinder clustering analysis.

We identified putatively orthologous genes in the OrthoFinder clusters for the genome and the genome-plus-transcriptome datasets. For both datasets the same pipeline was followed. Clusters with 1-to-1 orthology were retained. For clusters with median per-species membership equal to 1 and mean less than 2.5, a phylogenetic tree was inferred with RAxML (using the LG+G model). Each tree was visually inspected to acquire the largest possible monophyletic clan, and in-paralogues and spuriously included sequences were removed. Individual alignments of each locus were filtered using trimal [116] and then concatenated into a supermatrix using fastconcat [117]. The supermatrices were analysed with RAxML [65] with 100 ML bootstraps and PhyloBayes [118] (see Supplementary Table S12 for specifications). Trees were summarised in FigTree.

### DATABASING AND DATA AVAILABILITY

All raw data have been deposited in the relevant INSDC databases. An assembly with minor curation (nHd3.1) has been deposited at DDBJ/ENA/GenBank under the accession MTYJ00000000. All RNA-Seq data are uploaded to GEO and SRA under the accession IDs GSE94295 and SRP098585, and the PacBio raw reads and miRNA-Seq data into SRA under the accession IDs SRX2495681 and SRX2495676. Accession IDs for each individual sequence file are stated in Supplementary Table S1. We set up a dedicated Ensembl genome browser (version 85) [54] using the EasyImport pipeline [119] and imported the *H. dujardini* genome and annotations described in this paper and the new gene predictions for *R. varieornatus*. These data are available to browse, query and download at 
http://www.tardigrades.org.

**SOFTWARE USAGE AND DATA MANIPULATION**

We used open source software tools where available, as detailed in Supplementary Table S12. Custom scripts developed for the project are uploaded to https://github.com/abs-yy/Hypsibius_dujardini_manuscript. We used G-language Genome Analysis Environment [120, 121] for sequence manipulation.

## Acknowledgement

The authors thank Nozomi Abe and Yuki Takai for experimental support, and Brett T. Hannigan at DNAnexus for advises on Falcon assembly. *Chlorella vulgaris* used to feed the tardigrades was provided courtesy of Chlorella Industry Co., Ltd. This work was supported by KAKENHI Grant-in-Aid for Young Scientists (No.22681029) from the Japan Society for the Promotion of Science (JSPS), by a Grant for Basic Science Research Projects from The Sumitomo Foundation (No.140340), by research funds from the Yamagata Prefectural Government and Tsuruoka City, Japan, and in part by the Sequencing Grant Program by Tomy Digital Biology Co., Ltd. GK was funded by a BBSRC PhD studentship. DRL is funded by a James Hutton Institute/School of Biological Sciences University of Edinburgh studentship. SK is funded by BBSRC award BB/K020161/1. LS is funded by a Baillie Gifford Studentship, University of Edinburgh.

## Author Contributions

YY, GDK, DRL, LS, SK performed informatics analyses, DDH found the conditions for effective anhydrobiosis, KI sequenced small RNAs, MT managed the computational resources, GK, KA performed sequencing and assembly, TK, HK, AT, TK provided the *Ramazzottius varieornatus* genome prior to publication. All members participated in writing the manuscript. MB and KA supervised the project.

## Reference

1. Dunn CW, Hejnol A, Matus DQ, Pang K, Browne WE, Smith SA, et al. Broad phylogenomic sampling improves resolution of the animal tree of life. Nature. 2008;452(7188):745–9. doi: 10.1038/nature06614. PubMed PMID: 18322464.

2. Campbell LI, Rota-Stabelli O, Edgecombe GD, Marchioro T, Longhorn SJ, Telford MJ, et al. MicroRNAs and phylogenomics resolve the relationships of Tardigrada and suggest that velvet worms are the sister group of Arthropoda. Proc Natl Acad Sci USA. 2011;108(38):15920–4. doi: 10.1073/pnas. 1105499108. PubMed PMID: 21896763; PubMed Central PMCID: PMCPMC3179045.

3. Borner J, Rehm P, Schill RO, Ebersberger I, Burmester T. A transcriptome approach to ecdysozoan phylogeny. Mol Phylogenet Evol. 2014;80:79–87. doi: 10.1016/j.ympev.2014.08.001. PubMed PMID: 25124096.

4. Edgecombe GD. Arthropod phylogeny: an overview from the perspectives of morphology, molecular data and the fossil record. Arthropod Struct Dev. 2010;39(2-3):74–87. doi: 10.1016/j.asd.2009.10.002. PubMed PMID: 19854297.

5. Nielsen C. The triradiate sucking pharynx in animal phylogeny. Invertebrate Biology. 2013;132(1):1–13. doi: 10.1111/ivb.12010.

6. Degma P, Bertolani R, Guidetti R. Actual checklist of Tardigrada species 2016 [cited 2017 FEB 16]. 2016/12/15:[1–47. Available from: http://www.tardigrada.modena.unimo.it./miscellanea/ActualchecklistofTardigrada.pdf.

7. Clegg JS. Cryptobiosis--a peculiar state of biological organization. Comp Biochem Physiol B Biochem Mol Biol. 2001; 128(4):613–24. PubMed PMID: 11290443.

8. Horikawa DD, Cumbers J, Sakakibara I, Rogoff D, Leuko S, Harnoto R, et al. Analysis of DNA repair and protection in the Tardigrade Ramazzottius varieornatus and Hypsibius dujardini after exposure to UVC radiation. PLoS One. 2013;8(6):e64793. doi: 10.1371/journal.pone.0064793. PubMed PMID: 23762256; PubMed Central PMCID: PMCPMC3675078.

9. Altiero T, Guidetti R, Caselli V, Cesari M, Rebecchi L. Ultraviolet radiation tolerance in hydrated and desiccated eutardigrades. J Zool Syst Evol Res. 2011;49:104–10. doi: 10.1111/j.1439-0469.2010.00607.x. PubMed PMID: WOS:000289799800018.

10. Hengherr S, Worland MR, Reuner A, Brummer F, Schill RO. Freeze tolerance, supercooling points and ice formation: comparative studies on the subzero temperature survival of limno-terrestrial tardigrades. J Exp Biol. 2009;212(Pt 6):802–7. doi: 10.1242/jeb.025973. PubMed PMID: 19251996.

11. Jonsson KI, Harms-Ringdahl M, Torudd J. Radiation tolerance in the eutardigrade Richtersius coronifer. Int J Radiat Biol. 2005;81(9):649–56. doi: 10.1080/09553000500368453. PubMed PMID: 16368643.

12. May RM, Maria M, Gumard J. Action différentielle des rayons x et ultraviolets sur le tardigrade Macrobiotus areolatus, a l’état actif et desséché. Bull Biol Fr Belg. 1964;98:349–36719.

13. Persson D, Halberg KA, Jorgensen A, Ricci C, Mobjerg N, Kristensen RM. Extreme stress tolerance in tardigrades: surviving space conditions in low earth orbit. J Zool Syst Evol Res. 2011;49:90–7. doi: 10.1111/j.l43904692010.00605.x. PubMed PMID: WOS:000289799800016.

14. Ramløv H, Westh P. Cryptobiosis in the Eutardigrade Adorybiotus (Richtersius) coronifer: Tolerance to Alcohols, Temperature and de novo Protein Synthesis. Zool Anz. 2001;240(3-4):517–23. doi: 10.1078/00445231-00062.

15. Ono F, Mori Y, Takarabe K, Fujii A, Saigusa M, Matsushima Y, et al. Effect of ultra-high pressure on small animals, tardigrades and Artemia. Cogent Physics. 2016;3(1):ll67575. doi: 10.l080/23311940.2016.1167575.

16. Beltrán-Pardo EA, Jönsson I, Wojcik A, Haghdoost S, Bermúdez Cruz RM, Bernal Villegas JE. Sequence analysis of the DNA-repair gene rad51 in the tardigrades Milnesium cf. tardigradum, Hypsibius dujardini and Macrobiotus cf. harmsworthi. J Limnol. 2013;72(s1):80–91. doi: l0.408l/jlimnol.2013.s1.e10.

17. Mali B, Grohme MA, Forster F, Dandekar T, Schnolzer M, Reuter D, et al. Transcriptome survey of the anhydrobiotic tardigrade Milnesium tardigradum in comparison with Hypsibius dujardini and Richtersius coronifer. BMC Genomics. 2010;11(168):168. doi: 10.1186/1471-2164-11-168. PubMed PMID: 20226016; PubMed Central PMCID: PMCPMC2848246.

18. Gusev O, Suetsugu Y, Cornette R, Kawashima T, Logacheva MD, Kondrashov AS, et al. Comparative genome sequencing reveals genomic signature of extreme desiccation tolerance in the anhydrobiotic midge. Nat Commun. 2014;5(4784):1–9. doi: 10.1038/ncomms5784. PubMed PMID: 25216354; PubMed Central PMCID: PMC4175575.

19. Boothby TC, Tenlen JR, Smith FW, Wang JR, Patanella KA, Nishimura EO, et al. Evidence for extensive horizontal gene transfer from the draft genome of a tardigrade. Proc Natl Acad Sci U S A. 2015;112(52):15976–81. doi: 10.1073/pnas.1510461112. PubMed PMID: 26598659; PubMed Central PMCID: PMCPMC4702960.

20. Koutsovoulos G, Kumar S, Laetsch DR, Stevens L, Daub J, Conlon C, et al. No evidence for extensive horizontal gene transfer in the genome of the tardigrade Hypsibius dujardini. Proc Natl Acad Sci U S A. 2016;113(18):5053–8. doi: 10.1073/pnas.1600338113. PubMed PMID: 27035985; PubMed Central PMCID: PMCPMC4983863.

21. Arakawa K. No evidence for extensive horizontal gene transfer from the draft genome of a tardigrade. Proc Natl Acad Sci U S A. 2016;3(22):E3057. doi: 10.1073/pnas.1602711113. PubMed PMID: 27173901; PubMed Central PMCID: PMCPMC4896722.

22. Hashimoto T, Horikawa DD, Saito Y, Kuwahara H, Kozuka-Hata H, Shin IT, et al. Extremotolerant tardigrade genome and improved radiotolerance of human cultured cells by tardigrade-unique protein. Nat Commun. 2016;7:12808. doi: 10.1038/ncomms12808. PubMed PMID: 27649274; PubMed Central PMCID: PMCPMC5034306.

23. Kondo K, Kubo T, Kunieda T. Suggested Involvement of PPl/PP2A Activity and De Novo Gene Expression in Anhydrobiotic Survival in a Tardigrade, Hypsibius dujardini, by Chemical Genetic Approach. PLoS One. 2015;10(12):e0144803. doi: 10.1371/journal.pone.0144803. PubMed PMID: 26690982; PubMed Central PMCID: PMCPMC4686906.

24. Horikawa DD, Kunieda T, Abe W, Watanabe M, Nakahara Y, Yukuhiro F, et al. Establishment of a rearing system of the extremotolerant tardigrade Ramazzottius varieornatus: a new model animal for astrobiology. Astrobiology. 2008;8(3):549–56. doi: l0.l089/ast.2007.0139. PubMed PMID: 18554084.

25. Tenlen JR, McCaskill S, Goldstein B. RNA interference can be used to disrupt gene function in tardigrades. Dev Genes Evol. 2013;223(3):171–81. doi: 10.1007/s00427-012-0432-6. PubMed PMID: 23187800; PubMed Central PMCID: PMCPMC3600081.

26. Horikawa D. The tardigrade Ramazzottius varieornatus as a model of extremotolerant animals. J Jpn Soc Extremophiles. 2008;7.2(1):25–8. doi: 10.3118/jjse.7.2.25.

27. Rizzo AM, Negroni M, Altiero T, Montorfano G, Corsetto P, Berselli P, et al. Antioxidant defences in hydrated and desiccated states of the tardigrade Paramacrobiotus richtersi. Comp Biochem Physiol B Biochem Mol Biol. 2010;156(2):115–21. doi: 10.1016/j.cbpb.2010.02.009. PubMed PMID: 20206711.

28. Jonsson KI, Schill RO. Induction of Hsp70 by desiccation, ionising radiation and heat-shock in the eutardigrade Richtersius coronifer. Comp Biochem Physiol B Biochem Mol Biol. 2007;146(4):456–60. doi: 10.1016/j.cbpb.2006.10.111. PubMed PMID: 17261378.

29. Reuner A, Hengherr S, Mali B, Forster F, Arndt D, Reinhardt R, et al. Stress response in tardigrades: differential gene expression of molecular chaperones. Cell Stress Chaperones. 2010;15(4):423–30. doi: 10.1007/s12192-009-0158-1. PubMed PMID: 19943197; PubMed Central PMCID: PMCPMC3082643.

30. Schill RO, Steinbruck GH, Kohler HR. Stress gene (hsp70) sequences and quantitative expression in Milnesium tardigradum (Tardigrada) during active and cryptobiotic stages. J Exp Biol. 2004;207(Pt 10):1607–13. PubMed PMID: 15073193.

31. Hengherr S, Heyer AG, Kohler HR, Schill RO. Trehalose and anhydrobiosis in tardigrades--evidence for divergence in responses to dehydration. FEBS J. 2008;275(2):281–8. doi: 10.1111/j.1742- 4658.2007.06198.x. PubMed PMID: 18070104.

32. Moore DS, Hansen R, Hand SC. Liposomes with diverse compositions are protected during desiccation by LEA proteins from Artemia franciscana and trehalose. Biochim Biophys Acta. 2016;1858(1):104–15. doi: 10.1016/j.bbamem.2015.10.019. PubMed PMID: 26518519.

33. Yamaguchi A, Tanaka S, Yamaguchi S, Kuwahara H, Takamura C, Imajoh-Ohmi S, et al. Two novel heat-soluble protein families abundantly expressed in an anhydrobiotic tardigrade. PLoS One. 2012;7(8):e44209. doi: 10.1371/journal.pone.0044209. PubMed PMID: 22937162; PubMed Central PMCID: PMCPMC3429414.

34. Tanaka S, Tanaka J, Miwa Y, Horikawa DD, Katayama T, Arakawa K, et al. Novel mitochondria-targeted heat-soluble proteins identified in the anhydrobiotic Tardigrade improve osmotic tolerance of human cells. PLoS One. 2015;10(2):e0118272. doi: 10.1371/journal.pone.0118272. PubMed PMID: 25675104; PubMed Central PMCID: PMCPMC4326354.

35. Kikawada T, Saito A, Kanamori Y, Nakahara Y, Iwata K, Tanaka D, et al. Trehalose transporter 1, a facilitated and high-capacity trehalose transporter, allows exogenous trehalose uptake into cells. Proc Natl Acad Sci U S A. 2007; 104(28): 11585–90. doi: 10.1073/pnas.0702538104. PubMed PMID: 17606922; PubMed Central PMCID: PMCPMC1905927.

36. da Costa Morato Nery D, da Silva CG, Mariani D, Fernandes PN, Pereira MD, Panek AD, et al. The role of trehalose and its transporter in protection against reactive oxygen species. Biochim Biophys Acta. 2008; 1780(12):1408–11. doi: 10.1016/j.bbagen.2008.05.011. PubMed PMID: 18601980.

37. Madin KAC, Crowe JH. Anhydrobiosis in nematodes: Carbohydrate and lipid metabolism during dehydration. Journal of Experimental Zoology. 1975;193(3):335–42. doi: 10.1002/jez.1401930309.

38. Clegg JS. Metabolic studies of crytobiosis in encysted embryos of Artemia salina. Comparative Biochemistry and Physiology. 1967;20(3):801–9. doi: 10.1016/0010406(67)900540-0.

39. Bemm F, Weiss CL, Schultz J, Forster F. Genome of a tardigrade: Horizontal gene transfer or bacterial contamination? Proc Natl Acad Sci U S A. 2016;113(22):E3054–6. doi: 10.1073/pnas. 1525116113. PubMed PMID: 27173902; PubMed Central PMCID: PMCPMC4896698.

40. Delmont TO, Eren AM. Identifying contamination with advanced visualization and analysis practices: metagenomic approaches for eukaryotic genome assemblies. PeerJ. 2016;4:e1839. doi: 10.7717/peerj. 1839. PubMed PMID: 27069789; PubMed Central PMCID: PMCPMC4824900.

41. Boothby TC, Goldstein B. Reply to Bemm et al. and Arakawa: Identifying foreign genes in independent Hypsibius dujardini genome assemblies. Proc Natl Acad Sci U S A. 2016;113(22):E3058–61. doi: 10.1073/pnas. 1601149113. PubMed PMID: 27173900; PubMed Central PMCID: PMCPMC4896697.

42. Arakawa K, Yoshida Y, Tomita M. Genome sequencing of a single tardigrade *Hypsibius dujardini* individual. Sci Data. 2016;3:160063. doi: 10.1038/sdata.2016.63. PubMed PMID: 27529330; PubMed Central PMCID: PMCPMC4986543.

43. Chin CS, Peluso P, Sedlazeck FJ, Nattestad M, Concepcion GT, Clum A, et al. Phased diploid genome assembly with single-molecule real-time sequencing. Nat Methods. 2016;13(12):1050–4. doi: 10.1038/nmeth.4035. PubMed PMID: 27749838.

44. Kajitani R, Toshimoto K, Noguchi H, Toyoda A, Ogura Y, Okuno M, et al. Efficient de novo assembly of highly heterozygous genomes from whole-genome shotgun short reads. Genome Res. 2014;24(8):1384–95. doi: 10.1101/gr. 170720.113. PubMed PMID: 24755901; PubMed Central PMCID: PMCPMC4120091.

45. Gabriel WN, McNuff R, Patel SK, Gregory TR, Jeck WR, Jones CD, et al. The tardigrade Hypsibius dujardini, a new model for studying the evolution of development. Dev Biol. 2007;312(2):545–59. doi: 10.1016/j.ydbio.2007.09.055. PubMed PMID: 17996863.

46. Boetzer M, Pirovano W. SSPACE-LongRead: scaffolding bacterial draft genomes using long read sequence information. BMC Bioinformatics. 2014;15(211):211. doi: 10.1186/1471-2105-15-211. PubMed PMID: 24950923; PubMed Central PMCID: PMCPMC4076250.

47. Parra G, Bradnam K, Korf I. CEGMA: a pipeline to accurately annotate core genes in eukaryotic genomes. Bioinformatics. 2007;23(9):1061–7. doi: 10.1093/bioinformatics/btm071. PubMed PMID: 17332020.

48. Hoff KJ, Lange S, Lomsadze A, Borodovsky M, Stanke M. BRAKER1: Unsupervised RNA-Seq-Based Genome Annotation with GeneMark-ET and AUGUSTUS. Bioinformatics. 2016;32(5):767–9. doi: 10.1093/bioinformatics/btv661. PubMed PMID: 26559507.

49. Grabherr MG, Haas BJ, Yassour M, Levin JZ, Thompson DA, Amit I, et al. Full-length transcriptome assembly from RNA-Seq data without a reference genome. Nat Biotechnol. 2011; 29(7):644–52. doi: 10.1038/nbt. 1883. PubMed PMID: 21572440; PubMed Central PMCID: PMCPMC3571712.

50. Lagesen K, Hallin P, Rodland EA, Staerfeldt HH, Rognes T, Ussery DW. RNAmmer: consistent and rapid annotation of ribosomal RNA genes. Nucleic Acids Res. 2007;35(9):3100–8. doi: 10.1093/nar/gkm160. PubMed PMID: 17452365; PubMed Central PMCID: PMCPMC1888812.

51. Lowe TM, Eddy SR. tRNAscan-SE: a program for improved detection of transfer RNA genes in genomic sequence. Nucleic Acids Res. 1997;25(5):955–64. doi: 10.1093/nar/25.5.0955. PubMed PMID: 9023104; PubMed Central PMCID: PMCPMC146525.

52. Friedlander MR, Mackowiak SD, Li N, Chen W, Rajewsky N. miRDeep2 accurately identifies known and hundreds of novel microRNA genes in seven animal clades. Nucleic Acids Res. 2012;40(1):37–52. doi: 10.1093/nar/gkr688. PubMed PMID: 21911355; PubMed Central PMCID: PMCPMC3245920.

53. Kozomara A, Griffiths-Jones S. miRBase: annotating high confidence microRNAs using deep sequencing data. Nucleic Acids Res. 2014;42(Database issue):D68–73. doi: 10.1093/nar/gkt 1181. PubMed PMID: 24275495; PubMed Central PMCID: PMCPMC3965103.

54. Aken BL, Achuthan P, Akanni W, Amode MR, Bernsdorff F, Bhai J, et al. Ensembl 2017. Nucleic Acids Res. 2017;45(D1):D635–D42. doi: 10.1093/nar/gkw1 104. PubMed PMID: 27899575; PubMed Central PMCID: PMCPMC5210575.

55. Yates A, Beal K, Keenan S, McLaren W, Pignatelli M, Ritchie GR, et al. The Ensembl REST API: Ensembl Data for Any Language. Bioinformatics. 2015;31(1):143–5. doi: 10.1093/bioinformatics/btu613. PubMed PMID: 25236461; PubMed Central PMCID: PMCPMC4271150.

56. Hubley R, Finn RD, Clements J, Eddy SR, Jones TA, Bao W, et al. The Dfam database of repetitive DNA families. Nucleic Acids Res. 2016;44(D1):D81–9. doi: 10.1093/nar/gkv1272. PubMed PMID: 26612867; PubMed Central PMCID: PMCPMC4702899.

57. UniProt C. UniProt: a hub for protein information. Nucleic Acids Res. 2015;43(Database issue):D204–12. doi: 10.1093/nar/gku989. PubMed PMID: 25348405; PubMed Central PMCID: PMCPMC4384041.

58. Popendorf K, Tsuyoshi H, Osana Y, Sakakibara Y. Murasaki: a fast, parallelizable algorithm to find anchors from multiple genomes. PLoS One. 2010;5(9):e12651. doi: 10.1371/journal.pone.0012651. PubMed PMID: 20885980; PubMed Central PMCID: PMCPMC2945767.

59. Emms DM, Kelly S. OrthoFinder: solving fundamental biases in whole genome comparisons dramatically improves orthogroup inference accuracy. Genome Biol. 2015;16:157. doi: 10.1186/s13059-015-0721-2. PubMed PMID: 26243257; PubMed Central PMCID: PMCPMC4531804.

60. Laetsch DR. KinFin v0.8.1.2017. doi: 10.5281/zenodo.290589.

61. Keller O, Kollmar M, Stanke M, Waack S. A novel hybrid gene prediction method employing protein multiple sequence alignments. Bioinformatics. 2011;27(6):757–63. doi: 10.1093/bioinformatics/btr010. PubMed PMID: 21216780.

62. Boschetti C, Carr A, Crisp A, Eyres I, Wang-Koh Y, Lubzens E, et al. Biochemical diversification through foreign gene expression in bdelloid rotifers. PLoS Genet. 2012;8(11):e1003035. doi: 10.1371/journal.pgen. 1003035. PubMed PMID: 23166508; PubMed Central PMCID: PMCPMC3499245.

63. Buchfink B, Xie C, Huson DH. Fast and sensitive protein alignment using DIAMOND. Nature Methods. 2015;12(1):59–60. PubMed PMID: WOS:000347668600019.

64. Altschul SF, Madden TL, Schaffer AA, Zhang J, Zhang Z, Miller W, et al. Gapped BLAST and PSI - BLAST: a new generation of protein database search programs. Nucleic Acids Res. 1997;25(17):3389–402. PubMed PMID: 9254694; PubMed Central PMCID: PMCPMC146917.

65. Stamatakis A. RAxML version 8: a tool for phylogenetic analysis and post-analysis of large phylogenies. Bioinformatics. 2014;30(9):1312–3. doi: 10.1093/bioinformatics/btu033. PubMed PMID: 24451623; PubMed Central PMCID: PMCPMC3998144.

66. Oliveira RP, Porter Abate J, Dilks K, Landis J, Ashraf J, Murphy CT, et al. Condition-adapted stress and longevity gene regulation by Caenorhabditis elegans SKN-1/Nrf. Aging Cell. 2009;8(5):524–41. doi: 10.1111/j.1474-9726.2009.00501.x.. PubMed PMID: 19575768; PubMed Central PMCID: PMCPMC2776707.

67. de Rosa R, Grenier JK, Andreeva T, Cook CE, Adoutte A, Akam M, et al. Hox genes in brachiopods and priapulids and protostome evolution. Nature. 1999;399(6738):772–6. doi: 10.1038/21631. PubMed PMID: 10391241.

68. Grenier JK, Garber TL, Warren R, Whitington PM, Carroll S. Evolution of the entire arthropod Hox gene set predated the origin and radiation of the onychophoran/arthropod clade. Current Biology. 1997;7(8):547–53. doi: 10.1016/s0960-9822(06)00253-3.

69. Janssen R, Eriksson BJ, Tait NN, Budd GE. Onychophoran Hox genes and the evolution of arthropod Hox gene expression. Front Zool. 2014;11(1):22. doi: 10.1186/1742-9994-11-22. PubMed PMID: 24594097; PubMed Central PMCID: PMCPMC4015684.

70. Smith FW, Boothby TC, Giovannini I, Rebecchi L, Jockusch EL, Goldstein B. The Compact Body Plan of Tardigrades Evolved by the Loss of a Large Body Region. Curr Biol. 2016;26(2):224–9. doi: 10.1016/j.cub.2015.11.059. PubMed PMID: 26776737.

71. Aboobaker A, Blaxter M. Hox gene evolution in nematodes: novelty conserved. Current Opinion in Genetics & Development. 2003;13(6):593–8. doi: 10.1016/j.gde.2003.10.009.

72. Aboobaker AA, Blaxter ML. Hox Gene Loss during Dynamic Evolution of the Nematode Cluster. Current Biology. 2003;13(1):37–40. doi: 10.1016/s0960-9822(02)01399-4.

73. Aboobaker A, Blaxter M. The Nematode Story: Hox Gene Loss and Rapid Evolution. In: Deutsch JS, editor. Hox Genes: Studies from the 20th to the 21st Century. New York, NY: Springer New York; 2010. p. 101–10.

74. Mitreva M, Blaxter ML, Bird DM, McCarter JP. Comparative genomics of nematodes. Trends Genet. 2005;21(10):573–8. doi: 10.1016/j.tig.2005.08.003. PubMed PMID: 16099532.

75. Yoshinaga K, Yoshioka H, Kurosaki H, Hirasawa M, Uritani M, Hasegawa K. Protection by trehalose of DNA from radiation damage. Biosci Biotechnol Biochem. 1997;61(1):160–1. PubMed PMID: 9028044.

76. Herdeiro RS, Pereira MD, Panek AD, Eleutherio EC. Trehalose protects Saccharomyces cerevisiae from lipid peroxidation during oxidative stress. Biochim Biophys Acta. 2006;1760(3):340–6. doi: 10.1016/j.bbagen.2006.01.010. PubMed PMID: 16510250.

77. Grohme MA, Mali B, Welnicz W, Michel S, Schill RO, Frohme M. The Aquaporin Channel Repertoire of the Tardigrade Milnesium tardigradum. Bioinform Biol Insights. 2013;7: 153–65. doi: 10.4137/BB1.S11497. PubMed PMID: 23761966; PubMed Central PMCID: PMCPMC3666991.

78. Cornette R, Kikawada T. The induction of anhydrobiosis in the sleeping chironomid: current status of our knowledge. IUBMB Life. 2011;63(6):419–29. doi: 10.1002/iub.463. PubMed PMID: 21547992.

79. Gross V, Mayer G. Neural development in the tardigrade Hypsibius dujardini based on anti-acetylated alpha-tubulin immunolabeling. Evodevo. 2015;6:12. doi: 10.11 86/s13227-015-0008-4. PubMed PMID: 26052416; PubMed Central PMCID: PMCPMC4458024.

80. Martin C, Gross V, Pflüger H-J, Stevenson PA, Mayer G. Assessing segmental versus non-segmental features in the ventral nervous system of onychophorans (velvet worms). BMC Evolutionary Biology. 2017;17(1):3. doi: 10.1186/s12862-016-0853-3.

81. Sulston JE, Schierenberg E, White JG, Thomson JN. The embryonic cell lineage of the nematode Caenorhabditis elegans. Developmental Biology. 1983;100(1):64–119. doi: 10.1016/0012-1606(83)90201-4.

82. Andrews S. FastQC a quality-control tool for high-throughput sequence data. 2015 [cited 2015 May 21]. Available from: http://www.bioinformatics.babraham.ac.uk/projects/fastqc/.

83. Edgar RC. Search and clustering orders of magnitude faster than BLAST. Bioinformatics. 2010;26(19):2460–1. doi: 10.1093/bioinformatics/btq461. PubMed PMID: 20709691.

84. Bankevich A, Nurk S, Antipov D, Gurevich AA, Dvorkin M, Kulikov AS, et al. SPAdes: a new genome assembly algorithm and its applications to single-cell sequencing. J Comput Biol. 2012;19(5):455–77. doi: 10.1089/cmb.2012.0021. PubMed PMID: 22506599; PubMed Central PMCID: PMCPMC3342519.

85. Camacho C, Coulouris G, Avagyan V, Ma N, Papadopoulos J, Bealer K, et al. BLAST+: architecture and applications. BMC Bioinformatics. 2009;10:421. doi: 10.1186/1471-2105-10-42. PubMed PMID: 20003500; PubMed Central PMCID: PMCPMC2803857.

86. O'Leary NA, Wright MW, Brister JR, Ciufo S, Haddad D, McVeigh R, et al. Reference sequence (RefSeq) database at NCBI: current status, taxonomic expansion, and functional annotation. Nucleic Acids Res. 2016;44(D1):D733–45. doi: 10.1093/nar/gkv1189. PubMed PMID: 26553804; PubMed Central PMCID: PMCPMC4702849.

87. Kumar S, Jones M, Koutsovoulos G, Clarke M, Blaxter M. Blobology: exploring raw genome data for contaminants, symbionts and parasites using taxon-annotated GC-coverage plots. Front Genet. 2013;4:237. doi: 10.3389/fgene.2013.00237. PubMed PMID: 24348509; PubMed Central PMCID: PMCPMC3843372.

88. Langmead B, Salzberg SL. Fast gapped-read alignment with Bowtie 2. Nat Methods. 2012;9(4):357–9. doi: 10. 1038/nmeth. 1923. PubMed PMID: 22388286; PubMed Central PMCID: PMCPMC3322381.

89. English AC, Richards S, Han Y, Wang M, Vee V, Qu J, et al. Mind the gap: upgrading genomes with Pacific Biosciences RS long-read sequencing technology. PLoS One. 2012;7(11):e47768. doi: 10.1371/journal.pone.0047768. PubMed PMID: 23185243; PubMed Central PMCID: PMCPMC3504050.

90. Walker BJ, Abeel T, Shea T, Priest M, Abouelliel A, Sakthikumar S, et al. Pilon: an integrated tool for comprehensive microbial variant detection and genome assembly improvement. PLoS One. 2014;9(11):e112963. doi: 10.1371/journal.pone.0112963. PubMed PMID: 25409509; PubMed Central PMCID: PMCPMC4237348.

91. Kim D, Pertea G, Trapnell C, Pimentel H, Kelley R, Salzberg SL. TopHat2: accurate alignment of transcriptomes in the presence of insertions, deletions and gene fusions. Genome Biol. 2013;14(4):R36. doi: 10.1186/gb-2013-14-4-r36. PubMed PMID: 23618408; PubMed Central PMCID: PMCPMC4053844.

92. Borodovsky M, Lomsadze A. Eukaryotic gene prediction using GeneMark.hmm-E and GeneMark-ES. Curr Protoc Bioinformatics. 2011;Chapter 4:Unit 4 6 1–10. doi: 10.1002/0471250953.bi0406s35. PubMed PMID: 21901742; PubMed Central PMCID: PMCPMC3204378.

93. Li H, Durbin R. Fast and accurate short read alignment with Burrows-Wheeler transform. Bioinformatics. 2009;25(14):1754–60. doi: 10.1093/bioinformatics/btp324. PubMed PMID: 19451168; PubMed Central PMCID: PMCPMC2705234.

94. Li H, Handsaker B, Wysoker A, Fennell T, Ruan J, Homer N, et al. The Sequence Alignment/Map format and SAMtools. Bioinformatics. 2009;25(16):2078–9. doi: 10.1093/bioinformatics/btp352. PubMed PMID: 19505943; PubMed Central PMCID: PMCPMC2723002.

95. Okonechnikov K, Conesa A, Garcia-Alcalde F. Qualimap 2: advanced multi-sample quality control for high-throughput sequencing data. Bioinformatics. 2016;32(2):292–4. doi: 10.1093/bioinformatics/btv566. PubMed PMID: 26428292; PubMed Central PMCID: PMCPMC4708105.

96. Mistry J, Finn RD, Eddy SR, Bateman A, Punta M. Challenges in homology search: HMMER3 and convergent evolution of coiled-coil regions. Nucleic Acids Res. 2013;41(12):e121. doi: 10.1093/nar/gkt263. PubMed PMID: 23598997; PubMed Central PMCID: PMCPMC3695513.

97. Finn RD, Coggill P, Eberhardt RY, Eddy SR, Mistry J, Mitchell AL, et al. The Pfam protein families database: towards a more sustainable future. Nucleic Acids Res. 2016;44(D1):D279–85. doi: 10.1093/nar/gkv 1344. PubMed PMID: 26673716; PubMed Central PMCID: PMCPMC4702930.

98. Moriya Y, Itoh M, Okuda S, Yoshizawa AC, Kanehisa M. KAAS: an automatic genome annotation and pathway reconstruction server. Nucleic Acids Res. 2007;35(Web Server issue):W182–5. doi: 10.1093/nar/gkm321. PubMed PMID: 17526522; PubMed Central PMCID: PMCPMC1933193.

99. Goujon M, McWilliam H, Li W, Valentin F, Squizzato S, Paern J, et al. A new bioinformatics analysis tools framework at EMBL-EBI. Nucleic Acids Res. 2010;38(Web Server issue):W695–9. doi: 10.1093/nar/gkq313. PubMed PMID: 20439314; PubMed Central PMCID: PMCPMC2896090.

100. Price AL, Jones NC, Pevzner PA. De novo identification of repeat families in large genomes. Bioinformatics. 2005;21 Suppl 1:i351–8. doi: 10.1093/bioinformatics/bti1018. PubMed PMID: 15961478.

101. Smit A, Hubley R, Green P. RepeatMasker Open-4.0 2013-2015". Available from: http://www.repeatmasker.org.

102. Pearson WR, Lipman DJ. Improved tools for biological sequence comparison. Proc Natl Acad Sci U S A. 1988;85(8):2444–8. PubMed PMID: 3162770; PubMed Central PMCID: PMCPMC280013.

103. Katoh K, Standley DM. MAFFT multiple sequence alignment software version 7: improvements in performance and usability. Mol Biol Evol. 2013;30(4):772–80. doi: 10.1093/molbev/mst010. PubMed PMID: 23329690; PubMed Central PMCID: PMCPMC3603318.

104. Love MI, Huber W, Anders S. Moderated estimation of fold change and dispersion for RNA-seq data with DESeq2. Genome Biol. 2014;15(12):550. doi: 10.1186/s13059-014-0550-8. PubMed PMID: 25516281; PubMed Central PMCID: PMCPMC4302049.

105. Bray NL, Pimentel H, Melsted P, Pachter L. Near-optimal probabilistic RNA-seq quantification. Nat Biotechnol. 2016;34(5):525–7. doi: 10.1038/nbt.3519. PubMed PMID: 27043002.

106. Okuda S, Yamada T, Hamajima M, Itoh M, Katayama T, Bork P, et al. KEGG Atlas mapping for global analysis of metabolic pathways. Nucleic Acids Res. 2008;36(Web Server issue):W423–6. doi: 10.1093/nar/gkn282. PubMed PMID: 18477636; PubMed Central PMCID: PMCPMC2447737.

107. Darling AC, Mau B, Blattner FR, Perna NT. Mauve: multiple alignment of conserved genomic sequence with rearrangements. Genome Res. 2004; 14(7):1394–403. doi: 10.1101/gr.2289704. PubMed PMID: 15231754; PubMed Central PMCID: PMCPMC442156.

108. Price MN, Dehal PS, Arkin AP. FastTree 2--approximately maximum-likelihood trees for large alignments. PLoS One. 2010;5(3):e9490. doi: 10.1371/journal.pone.0009490. PubMed PMID: 20224823; PubMed Central PMCID: PMCPMC2835736.

109. Rambaut A. FigTree. 2016.

110. Howe KL, Bolt BJ, Shafie M, Kersey P, Berriman M. WormBase ParaSite-a comprehensive resource for helminth genomics. Mol Biochem Parasitol. 2016. doi: 10.1016/j.molbiopara.2016.11.005. PubMed PMID: 27899279.

111. Howe KL, Bolt BJ, Cain S, Chan J, Chen WJ, Davis P, et al. WormBase 2016: expanding to enable helminth genomic research. Nucleic Acids Res. 2016;44(D1):D774–80. doi: 10.1093/nar/gkv1217. PubMed PMID: 26578572; PubMed Central PMCID: PMCPMC4702863.

112. Sievers F, Wilm A, Dineen D, Gibson TJ, Karplus K, Li W, et al. Fast, scalable generation of high-quality protein multiple sequence alignments using Clustal Omega. Mol Syst Biol. 2011;7(539):539. doi: 10.1038/msb.2011.75. PubMed PMID: 21988835; PubMed Central PMCID: PMCPMC326l699.

113. Huang X. CAP3: A DNA Sequence Assembly Program. Genome Research. 1999;9(9):868–77. doi: 10.1101/gr.9.9.868.

114. Jiang H, Lei R, Ding SW, Zhu S. Skewer: a fast and accurate adapter trimmer for next-generation sequencing paired-end reads. BMC Bioinformatics. 2014;15:182. doi: 10.1186/1471-2105-15-182. PubMed PMID: 24925680; PubMed Central PMCID: PMCPMC4074385.

115. Haas BJ, Papanicolaou A. TransDecoder (Find Coding Regions Within Transcripts).

116. Capella-Gutierrez S, Silla-Martinez JM, Gabaldon T. trimAl: a tool for automated alignment trimming in large-scale phylogenetic analyses. Bioinformatics. 2009;25(15):1972–3. doi: 10.1093/bioinformatics/btp348. PubMed PMID: 19505945; PubMed Central PMCID: PMCPMC27I2344.

117. Kuck P, Longo GC. FASconCAT-G: extensive functions for multiple sequence alignment preparations concerning phylogenetic studies. Front Zool. 2014;11(1):81. doi: 10.1186/s12983-014-0081-x. PubMed PMID: 25426157; PubMed Central PMCID: PMCPMC4243772.

118. Lartillot N, Philippe H. A Bayesian mixture model for across-site heterogeneities in the amino-acid replacement process. Mol Biol Evol. 2004;21(6):1095–109. doi: 10.1093/mo1bev/msh112. PubMed PMID: 15014145.

119. Challis RJ, Kumar S, Stevens L, Blaxter M. EasyMirror and EasyImport: Simplifying the setup of a custom Ensembl database and Webserver for any species. PeerJ Preprints. 2016;4(e2401v1). doi: 10.7287/peerj.preprints.2401v1.

120. Arakawa K, Mori K, Ikeda K, Matsuzaki T, Kobayashi Y, Tomita M. G-language Genome Analysis Environment: a workbench for nucleotide sequence data mining. Bioinformatics. 2003;19(2):305–6. PubMed PMID: I2538262.

121. Arakawa K, Tomita M. G-language system as a platform for large-scale analysis of high-throughput omics data. J Pestic Sci. 2006;31(3):282–8. doi: I0.l584/jpestics.31.282. PubMed PMID: WOS:0002401 14400006.

